# A mutation rate model at the basepair resolution identifies the mutagenic effect of Polymerase III transcription

**DOI:** 10.1101/2022.08.20.504670

**Authors:** Vladimir Seplyarskiy, Daniel J. Lee, Evan M. Koch, Joshua S. Lichtman, Harding H. Luan, Shamil R. Sunyaev

## Abstract

*De novo* mutations occur with substantially different rates depending on genomic location, sequence context and DNA strand^1–4^. The success of many human genetics techniques, especially when applied to large population sequencing datasets with numerous recurrent mutations^5–7^, depends strongly on assumptions about the local mutation rate. Such techniques include estimation of selection intensity^8^, inference of demographic history^9^, and mapping of rare disease genes^10^. Here, we present Roulette, a genome-wide mutation rate model at the basepair resolution that incorporates known determinants of local mutation rate (http://genetics.bwh.harvard.edu/downloads/Vova/Roulette/). Roulette is shown to be more accurate than existing models^1,6^. Roulette has sufficient resolution at high mutation rate sites to model allele frequencies under recurrent mutation. We use Roulette to refine estimates of population growth within Europe by incorporating the full range of human mutation rates. The analysis of significant deviations from the model predictions revealed a 10-fold increase in mutation rate in nearly all genes transcribed by Polymerase III, suggesting a new mutagenic mechanism. We also detected an elevated mutation rate within transcription factor binding sites restricted to sites actively utilized in testis and residing in promoters.

---

The human single nucleotide mutation rate varies along the genome at different scales^4,11,12^. Some of this variation is explained by the combination of mutation type and immediately adjacent nucleotides, conceptualized as the mutation spectra^6,13^. The CpG di-nucleotide context induces by far the largest spectrum effect because of the strongly mutagenic effect of methylation at cytosines followed by guanine^14^. Previous studies demonstrated that the extended sequence context, well beyond the two adjacent bases, exerts an additional effect on mutation rates^1,15–17^. Mutation spectra also vary along the genome, indicating that rate differences are not fully explained by the surrounding DNA sequence^4,18^. Some of this variability tracks DNA properties like replication timing and gene expression^4^. Other effects, such as spikes of multinucleotide mutations in oocytes, lack obvious epigenetic correlates^4,19,20^. In addition to regional variation, the rates of many mutation types depend on the DNA strand^4,21–23^. Transcription alters the mutation spectra between transcribed and non-transcribed strands, while replication leads to differences between leading and lagging strands.

We developed “Roulette” a mutation rate model that incorporates these factors and more (see Methods). Each nucleotide has three potential mutations, and we hereafter refer to each of these potential mutations as a site. The extended sequence context is included by estimating the effect of the 6 upstream and 6 downstream nucleotides adjacent to each site (Figure 1a). Due to sparsity, it is impossible to accurately estimate the effect of each unique 12-nucleotide context. To account for this, we estimated the effect of the central pentamer (two nucleotides on either side) separately from the individual effects of the 8 more distant nucleotides, which are included as covariates (Figure 1a,b). For epigenomic features, Roulette incorporates methylation level (for both CpG transitions and CpG transversions), transcription direction, gene expression level in testis (for sites within gene bodies), and quantitative estimates of replication direction (Figure 1a,c). The incorporation of transcription and replication directions makes the model strand-dependent with unequal rates for mutations of the same type on the two DNA strands. To our knowledge, strand-dependency has not been incorporated into existing context-dependent and regional models^1,6,24^ of germline mutation rate, but was incorporated in the context of cancer mutagenesis^25^. The Roulette model accounts for local mutation rate variation by including the observed mutability of each tri-nucleotide context in 50KB windows (Figure 1d). Known epigenetic factors contribute to the regional variation of mutation rate at this scale, but the SNV density is a more direct proxy to the local mutation rates than noisy epigenetic tracks. This approach also has a benefit of accounting for the regional variation unexplained by existing epigenetic features ^17,18^ (Supplementary Figure 1-3). Some DNA repair pathways act differently in intergenic regions, gene bodies and promoters^26,27^. We fit separate statistical models for each genomic compartment and each pentamer, thereby allowing the effects of covariates to vary independently among compartments and pentamers (64118 models total, see Methods). SNV probabilities were modeled using logistic regression. We fit models with pairwise interactions (115 parameters) and without pairwise interactions (25 parameters) between all covariates and selected the best performing model using cross-validation based on a 50/50 test/train split (see Methods). Two simpler models were also analyzed to prevent overfitting in pentamer-compartment pairs with too few mutations. Finally, we grouped the predicted rates into 100 bins because discrete mutation rate classes facilitate many applications such as analyses of allele frequency distributions.

**Figure 1.**
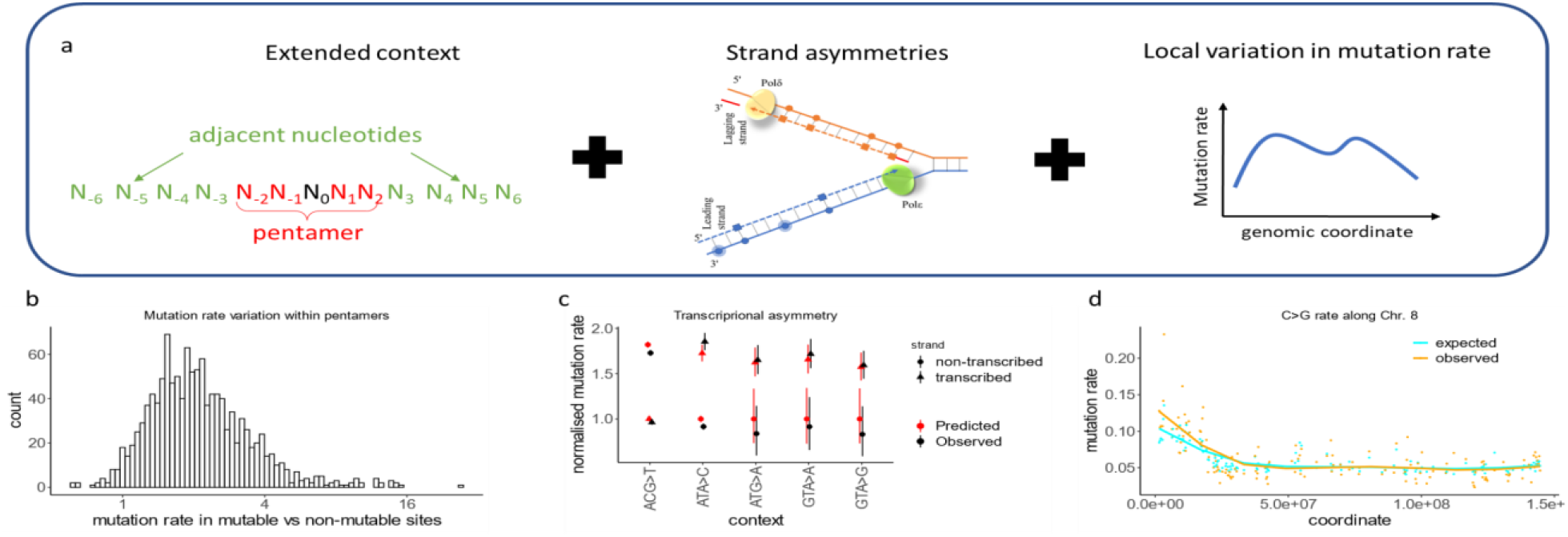
Roulette accounts for extended nucleotide context, strand asymmetries and local variation in mutation rate. a) Roulette is implemented as logistic regression with pairwise interactions (see Methods). For each pentamer, we model the effect of eight surrounding nucleotides (left), strand specific information (middle), and context-specific variation along the genome (right). b) Ratio of observed *de novo* mutation rates between the Roulette predicted most and least mutable deciles for each pentamer shows large variation unexplained by the pentamer context alone. c) Effect of transcriptional asymmetry on the rate of rare synonymous SNVs in the genes with high expression in testis (top quartile). Mutation rate is relative to the least mutable strand. d) Spike of the density of rare synonymous SNVs on the left arm of chromosome 8. This region is known to be affected by increased maternal mutagenesis^4,17,23,24^.

It is impossible to fit parameter-rich mutation models to currently available *de novo* mutation datasets because of data sparsity. To train Roulette, we collected all non-coding SNVs with frequency below 0.001 from gnomAD v3 whole genomes^6^ (524M rare SNVs total). The derived allele of these rare SNVs correspond to mutational events. We assume that rare alleles are always derived (as opposed to ancestral); simulations suggest that this is violated at most for one in 33,000 SNVs (see Methods). The distribution of very rare non-coding SNVs along the genome is primarily driven by mutation rate differences, with the effects of biased gene conversion, direct and background selection being negligible^1^.

Due to the sample size of contemporary human sequencing data, many rare SNVs represent recurrent mutations that have occurred multiple times in the genealogical history of the sequenced cohort. Because Roulette only fits the density of monomorphic sites, we transformed SNV probabilities to the mutation rate scale by assuming the probability a site remains monomorphic is given by the zero class of the Poisson distribution for the expected number of variants per site. The expected number of variants is proportional to mutation rate and the overall coalescent depth. We assume that the coalescent depth is approximately constant for a very large sample from a growing population (see Methods).

After estimation and rescaling, we found that Roulette captures expected genomic mutation rate variation when applied to synonymous sites not used in the model training. For instance, nearly two-fold rate differences between the transcribed and non-transcribed strands are predicted accurately (Figure 1c). Despite not using replication timing, histone modifications, or recombination rate^28^ as covariates, the direct inclusion of the regional variation is able to capture associations between mutability and these epigenetic factors (Supplementary Figure 2). The importance of this regional correction is illustrated by DNA segments that are hypermutable in oocytes (sometimes called regions of maternal mutagenesis)^19,20,29,30^. Maternal mutagenesis is responsible for a localized increase in C>G mutations on the left arm of the chromosome 8 (Figure 1d, Supplementary Figure 3).

As a second point of validation, we tested whether Roulette estimates resolve the old riddle of “cryptic variation.” Early comparative genomics literature^31–33^ observed that the frequency of triallelic SNVs is higher than expected based on the probability of pairs of biallelic SNVs assuming independence and a three nucleotide mutational model. Roulette accurately predicts the probability of triallelic SNVs (Supplementary Figure 4), suggesting that previous observations of “cryptic variation” reflected residual mutation rate variance in earlier models associated with extended nucleotide context and local genomic factors.

We compared Roulette with two existing mutation rate models to further validate its performance. Karczewski et al. (2020)^6^ which used trinucleotide context and methylation levels to estimate rates for gnomAD v2, and Carlson et al. (2018)^1^ which used heptamer context along with several epigenetic features including methylation levels for the BRIDGES study (see Supplementary Table 1 for a more detailed description of model differences). We re-fit the model from Karczewski et al. (2020)^6^ on gnomAD v3, and we used the publicly available estimates from Carlson et al. (2018). We hereafter refer to these models as gnomAD and Carlson.

While previous studies evaluated goodness of fit of mutation rate models^1^, none to our knowledge have attempted to estimate the remaining residual variance. We used two novel site-by-site metrics to analyze each model’s ability to predict the rate and location of observed SNVs from separate datasets. The first metric is an adjusted version of Nagelkerke’s pseudo-R^2^ for logistic models^34^ that measures the residual variance between the observed and expected likelihood given the inherently stochastic nature of mutational processes. Our Pseudo-R^2^ assumes that there is no variance among sites with the same predicted mutation rate, so that errors result solely from misclassification among mutation rate bins. We therefore developed a second per-site metric that estimates this additional variance within bins using observations of multiple mutations occurring at the same site. We compare the rate of *de novo* mutations at sites where an SNV was observed to the *de novo* rate at sites without SNVs. If the mutation rates corresponding to each bin are estimated without error, the *de novo* mutation rate in both groups should be equal. This SNV-conditional method uses the difference in *de novo* rates, depending on whether an SNV is observed or not, to estimate the within-bin variance. Both methods necessarily require assumptions about the true distribution of mutation rates. For pseudo-R^2^, we assume that the full distribution is well-captured by the model even if per-site estimates are subject to error, and for the SNV-conditional method, we assume that the distribution of true mutation rates within each bin is log-normal.

We first compared the models using synonymous variants from the gnomAD v2 whole exome dataset (∼125K individuals and ∼1.9M synonymous SNVs). Since Roulette was trained on non-coding variants only, synonymous variants are an independent dataset. Roulette predicts the rate of synonymous SNVs with higher accuracy than the Carlson and gnomAD models, reaching a pseudo-R^2^ of 0.86 compared to 0.81 and 0.78 respectively (Figure 2a). Next, we estimated pseudo-R^2^ in a UK Biobank whole genome sequencing dataset (200K individuals) for both synonymous (0.88, 0.83, 0.80) and non-coding sites (0.99, 0.94, 0.83) (Figure 2a, Supplementary Figure 5). Performances increased for non-coding likely because these were used in model training. Roulette’s Pseudo-R^2^ was also the highest for *de novo* synonymous mutations compiled from three independent trio-sequencing studies (41,816 trios and 2,759 *de novo* synonymous mutations; Pseudo-R^2^: 0.93, 0.87, 0.85)^29,35,36^.We assessed Roulette’s performance relative to the other mutational models using bootstrap samples of synonymous sites (see Methods) and showed that Roulette provides similar improvements across all validation sets (p<0.001; Figure 2b).

**Figure 2.**
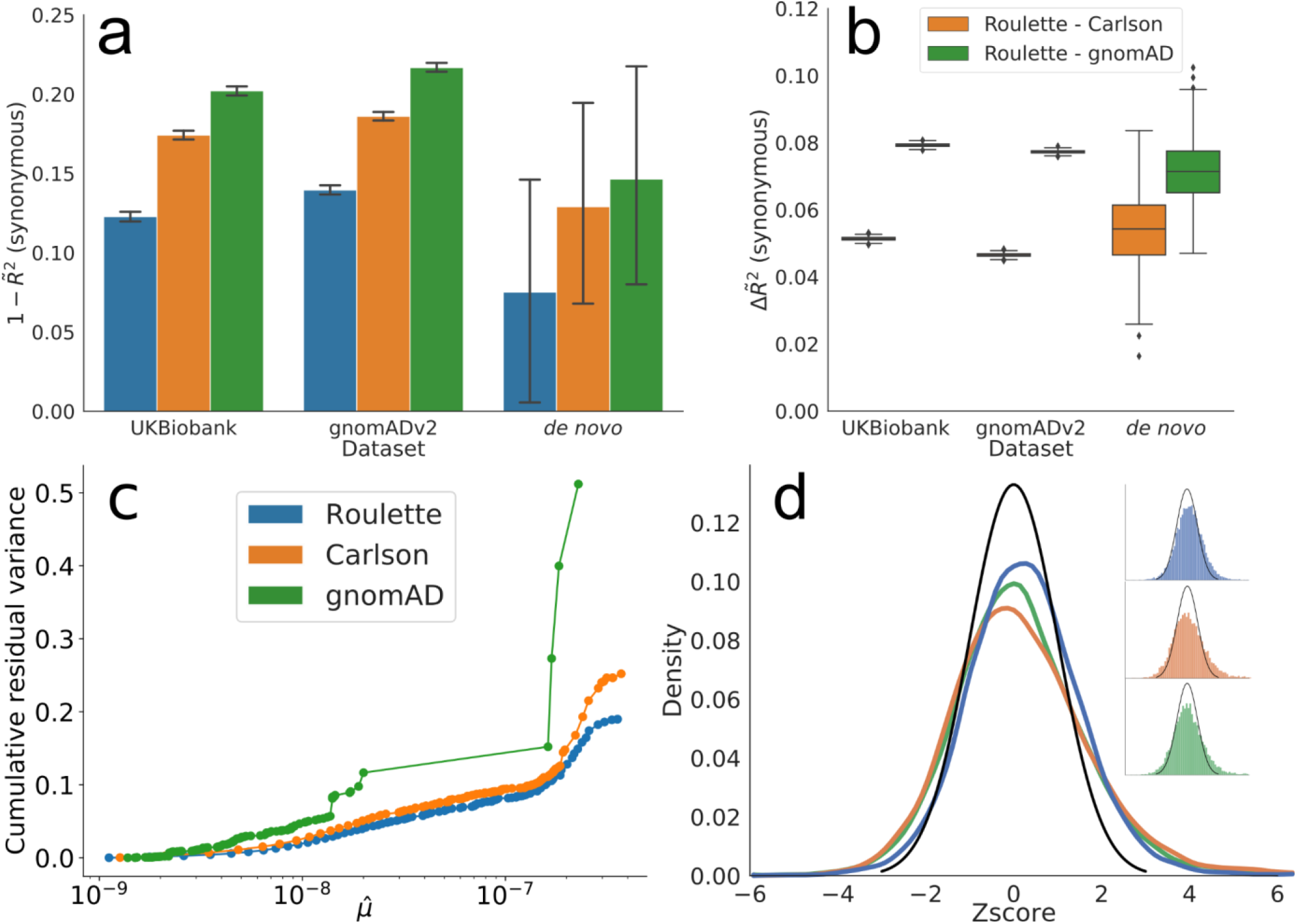
Roulette outperforms existing mutational models, under both per-gene and per-site metrics. a) 1 – pseudo-R^2^ of the three mutational models on synonymous variants observed in population sequencing data (gnomAD v2.1.1 and UK Biobank) and *de novo* mutation datasets^18,27,28^. A pseudo-R^2^ of 0 is equivalent to using genome-wide mean mutation rate for every site. A pseudo-R^2^ of 1 is the best per-site mutation rate estimate we can achieve, under the constraint that the mutation rates of synonymous sites follow the predicted genome-wide distribution. Error bars represent 95% confidence intervals estimated by bootstrap samples of synonymous sites. b) Difference in pseudo-R^2^ between Roulette and the two other models. The difference was calculated over each bootstrapped sample and whiskers represent estimated 95% confidence intervals. c) The estimated cumulative residual variance for the Carlson, gnomAD and Roulette models after binning mutation rate estimates. Within-bin variance is scaled by the total variance estimated for Roulette. The x-axis gives the estimated mean in each mutation rate bin scaled to the observed per-generation *de novo* rate observed in trio data. d) Error distributions on the Z-scale for predicted counts of synonymous mutations within genes in gnomAD v2. The standard normal density is shown in black to provide a reference for the expected error distribution if mutation rates were known without error.

As expected in the presence of residual mutation rate variation, sites that harbor SNVs in gnomAD had an excess of *de novo* mutations even when predicted mutation rates were within the same bin. The mean excess was 34% within Roulette bins, 47% within Carlson bins, and 94% within gnomAD bins (Supplementary Figure 6). These result in estimated residual variances of 19%, 25%, and 51% for the Roulette, Carlson, and gnomAD models (Figure 2c). While overall residual variances are larger for the SNV-conditional method, Roulette still explains around 5% more of the variance in human mutation rates than the Carlson model.

Many population genetics applications rely on aggregated mutation rate estimates by gene or within a genomic window. We evaluated the relevance of Roulette for these applications by aggregating synonymous sites by gene for gnomAD v2 and predicting the number of SNVs. Aggregate estimates generated using Roulette are more accurate than those for gnomAD or Carlson (Figure 2d). There are 1758 genes with a Z-score greater than 2 or less than -2 for Roulette rates, substantially fewer outlier genes than 2468 for Carlson or 2295 for the gnomAD model. An area of population genetics inference with important applications in human disease genetics is the estimation of selective constraints for protein truncating variants (PTVs). All methods to infer strong selection rely on estimates of local mutation rate. We recomputed estimates of two measures of strong heterozygous selection, ***s***_***het***_ and LOEUF^6,8^, using Roulette mutation rates. The new ***s***_***het***_ estimates (available at http://genetics.bwh.harvard.edu/genescores/selection.html) showed a slight but statistically significant improvement (p<0.001) in detection of autosomal dominant disease genes annotated in DDG2P (Supplementary Table 2), while updated LOEUF estimates showed no significant change.

We next investigated the utility of precise mutation rate estimates for the inference of demographic history (specifically, historical changes in effective population size) from the site frequency spectrum (SFS, the number of observed alleles at each frequency)^9,37^. Most studies rely on the “infinite number of sites” model which assumes that single mutation events contribute to each segregating site^38^. Under this model and neutrality, the relative distribution of allele frequencies only depends on genealogies and is independent of mutation rate, while the overall level of variation is linearly dependent on mutation rate. The presence of recent recurrent mutations breaks the key assumption of the infinite sites model and induces a dependency between the shape of site frequency spectrum and mutation rate^5,7,39^ (Figure 3a, Supplementary Figure 7). Using a set of SFS curves at different mutation rates can increase power and reduce biases due to recurrent mutations.

**Figure 3.**
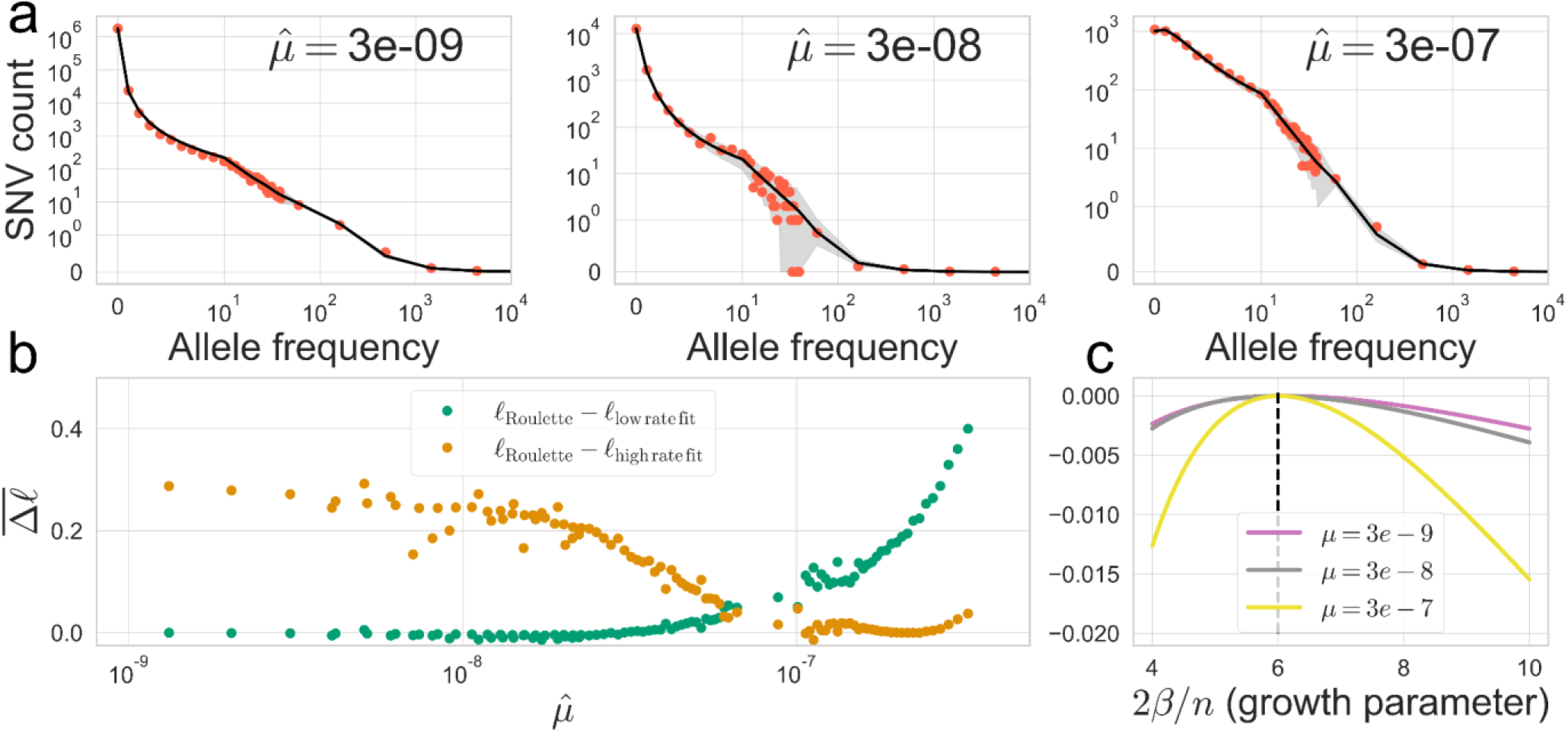
Accurate per-site mutation rate estimates improve population genetics inference. a) Estimated demographic history fits the SFS with mutation rate bins at different orders of magnitude. Red dots show the observed SFS at synonymous sites in gnomAD and black lines show the expected SFS under the inferred demographic model. Shaded areas correspond to 95% binomial confidence intervals. The observed SFS (red dots) shows the observed numbers of SNVs at allele counts 0-40. For more common alleles with counts above 40, red dots show numbers of SNVs for logarithmically (base 3) spaced bins. Allele counts are out of a total sample size of about 57K non-Finnish European individuals. b) Roulette bins improve fits to the shape of the SFS compared to demographic model predictions scaled to either low (1e-09 – 3.3e-09) or high rate (1e-07 – 3e-07) bins. Average log-likelihoods (per-SNV) are higher for Roulette after subtracting one to account for the additional parameter used to refit the mutation rate within each bin. Roulette improves over the model trained on sites with low mutation rate (mostly non-recurrent sites) because recurrent mutations change the shape of the SFS. It also improves over the high-rate model as one moves away from the mean mutation rate within the high-rate bin. c) High mutation rate SNVs are more informative about population growth parameters. The expected per-SNV log-likelihood relative to the maximum is shown using rare SNVs (1-40 allele counts). The compound population growth-rate / sample size parameter was chosen to approximate the observed synonymous SFS in gnomAD v2.

We re-fit a model of European demographic history^9^ to evaluate the ability of Roulette to model the shape of the SFS across the range of mutation rates. We used simulations that allowed for recurrent mutations^40^ to fit to the whole range of mutation rates. The inclusion of high mutation rate sites is meaningful because these are more informative per-variant about population growth than low-rate sites (Figure 3c). The demographic model allows for faster-than-exponential growth in the recent past, and we updated the acceleration parameter from 1.120 to 1.122 and the initial growth rate from 0.0050 to 0.0057 with a final population size estimated at 8.1 million compared to 2.5 million. This model fits the shape of the SFS well even as the mutation rate becomes large enough that recurrent mutation substantially skews to shape towards less rare variants^5,7,39^ (Figure 3a). The fine-scale mutation rate bins defined by Roulette provide a much better fit to the SFS shape than can be achieved by dividing mutations into only two bins, one for low rates and one for high (Figure 3b). This is due to sufficient recurrent mutation within the low-rate bin and sufficient rate variation within the high-rate bin to make single-rate summaries inadequate to capture the shape of the SFS. While one solution is to filter high mutation rates sites so that the infinite sites assumption remains reasonable, this removes sites that are more informative on a per-variant level (Figure 3c). This utility extends to selection inference where it is possible to identify individual strongly constrained sites when mutation rates are in the neighborhood of 1e-07 per generation^41^.

While much of mutation rate variation is adequately captured by Roulette (Figure 1, 2) including various epigenetically active sites like enhancers and promoters (Supplementary Figure 8), strong local deviations can be used to identify new mutagenic mechanisms in humans. Regional variation in mutation rates and spectra have previously been characterized and biologically interpreted at scales exceeding 10kb.^4,28^. However, many mutagenic mechanisms arise due to epigenetic factors acting at much shorter scales. Data sparsity prevents the application of unsupervised statistical techniques to characterize variation at short scales^4^. We analyzed extreme deviations from Roulette predictions at the 100bp scale genome-wide (Figure 4a). The choice of the scale was determined by the need to balance resolution and statistical power.

**Figure 4.**
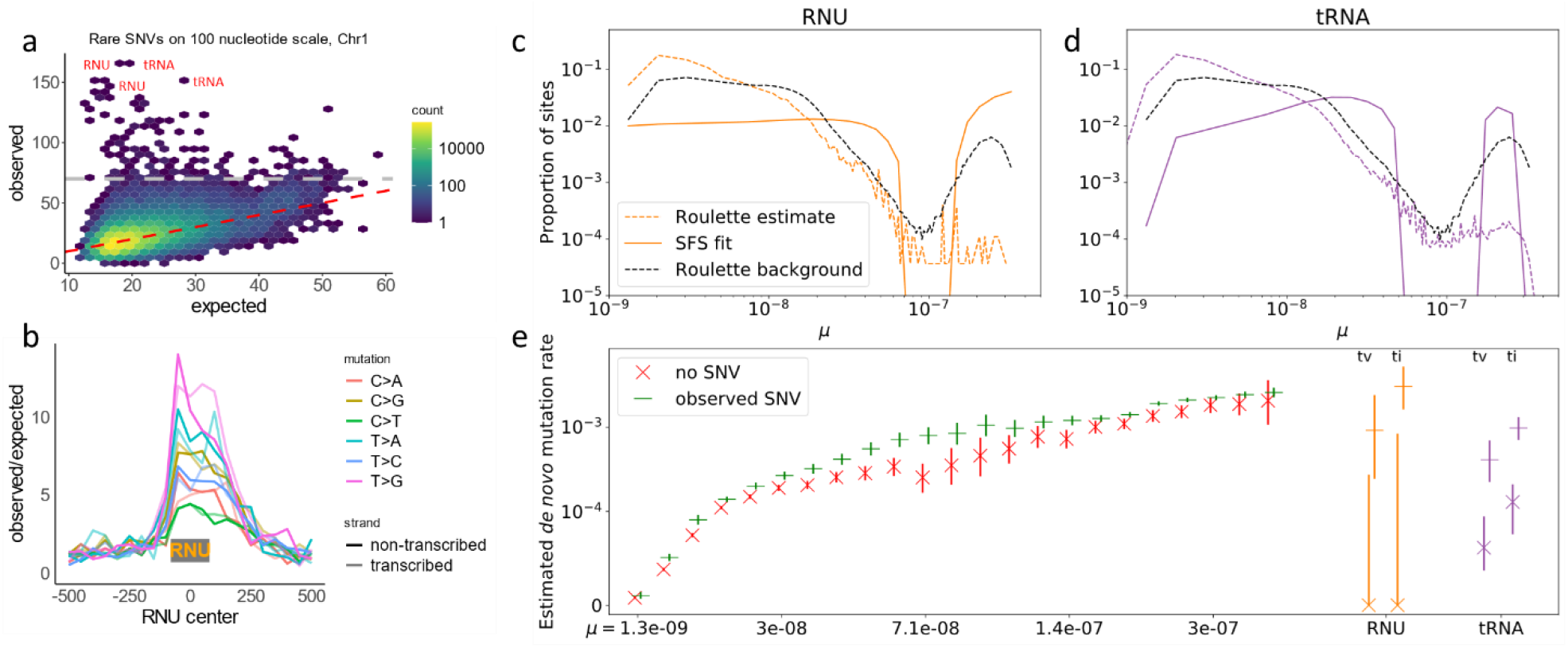
Polymerase III transcripts and transcription binding sites are mutational hotspots. a) Number of rare SNVs in 100 nucleotide non-overlapping windows. Expectation is calculated with Roulette. While mutation counts in most regions show minor deviations from the prediction, a few loci have much higher mutation rates (>70 SNVs, above the gray line). These loci are heavily enriched with Polymerase III transcripts. b) Mutation rate at and around small nuclear RNAs (RNU); the median RNU size is depicted as a gray rectangle. The mutation rate distributions for observed SNVs in c) RNU and d) tRNA genes was estimated by fitting the SFS in these genes as a mixture of SFS shapes observed in Roulette bins. Fits are compared to the original Roulette estimates and to the background rate distribution. e) SFS-based mutation rate predictions are validated by estimating the *de novo* rate for mutations with and without observed SNVs in gnomAD v3. Mutations are separated into transversions and transitions.

While many likely represent unfiltered sequencing and mapping artifacts, the most striking observation is that 25.6% of 100bp genomic windows with extremely high SNV counts unexplained by the Roulette features lie within RNA genes transcribed by polymerase III (Pol III). These outlier windows contain multiallelic variants and overall harbor over 70 SNVs per 100bp, while some windows have more than 100 SNVs. The two most prominent gene classes transcribed by Pol III are tRNA and small nuclear RNA genes (RNU) (Figure 4a, b, Supplementary Figure 9,10). Analyses of allelic imbalance and other sequence quality metrics suggested that these are true SNVs rather than sequencing artifacts (Supplementary Figure 11). We also found that the number of *de novo* mutations increases with paternal age, as expected for real germline mutations (Supplementary Figure 12). Elevated mutation rates in tRNA genes were recently noted by a comparative genomics study^42^, although the magnitude of the effect was likely underestimated by not accounting for recurrent mutations. Similarly, while we observe a 7-fold increase in SNV rate in RNU genes (Figure 4a,b), we expect that recurrent mutation means this is an underestimation of the true hypermutability. Indeed, we observed that *de novo* mutations in parent-child trio sequencing studies were detected at a 32-fold (19-50, 95% Poisson CI) higher rate in RNU genes.

To validate the link between Pol III transcription and elevated mutation rate, we compared mutation rates between active RNU genes and pseudogenes. The increased mutation rate is almost exclusively limited to active genes, suggesting that active transcription rather than genomic location or sequence context is responsible (Supplementary Figure 13a-c). The few exceptions are pseudogenes that show H3K27ac chromatin marks associated with active transcription (Supplementary Figure 13a,b), suggesting that apparent hypermutable RNU pseudogenes are misannotated active genes. The association between Pol III transcription and high SNV density extends to all other classes of non-coding RNAs (Supplementary Figure 13d) but not to SINE repeats (Supplementary Figure 13f), which may also be transcribed by Pol III^43^.

We next sought to further characterize the elevated mutation rates in tRNA and RNU genes. To this end, we developed a statistical model to estimate the distribution of mutation rates among observed SNVs (Supplementary Note 1) by taking advantage of the fact that recurrent mutations induce a shape-dependency of the SFS on mutation rate^5,7,39^. We observed that SFS in tRNAs and RNUs have the expected shift away from very rare variants, though milder than observed for variants at the top range of Roulette estimates (Supplementary Figure 14). By modeling SFS for a mixture of variants with different mutation rates, we estimated that mutation rate within Pol III transcripts is highly variable. Both RNU and tRNA genes have a large fraction of highly mutable sites, with mutability greatly exceeding the Roulette predictions (Figure 4c,d, Supplementary Figure 14). To validate the SFS-based predictions we calculated *de novo* mutation rates for RNU and tRNA sites, conditioning on the presence/absence of SNVs as well as transition/transversion status. There is a stark difference between estimated *de novo* mutation rates in polymorphic and monomorphic positions (Figure 4e), consistent with a high heterogeneity of mutation rate within RNU and tRNA genes SFS-based analysis and the analysis of *de novo* mutations in polymorphic sites estimated that the rate of RNU transitions in highly mutable sites is higher than for any Roulette bin (Figure 4c,e). RNU genes contain some of the most mutable positions in the human genome. The high mutation rate in Pol III transcripts masks the effect of purifying selection and leads to an unrealistic selection inference^44,45^.

There are multiple, not necessarily exclusive explanations as to why Pol III transcription is strongly mutagenic. First, unlike RNA polymerase II (Pol II), Pol III does not have the ability to recruit transcription coupled repair (TCR). However, TCR only removes mutations on one of the two strands and thus cannot reduce the mutation rate by more than a half. TCR alone is therefore insufficient to explain the observed 32-fold effect. Second, transcription associated mutagenesis (TAM), a well-described phenomenon in yeasts^46^, is attributed primarily to ribonucleotide incorporation into DNA during transcription. A third possibility could involve an as-of-yet uncharacterized transcription-associated mechanism specific to Pol III, because the biological machinery transcribing Pol III-dependent genes differs substantially from the machinery for Pol II-dependent genes^47^.

Interestingly, it was recently shown that damage-induced mutations can accumulate on the non-transcribed strand outside of replication^4,48^. However, this mechanism creates a very strong mutational asymmetry that is absent for Pol III transcripts. Finally, transcription initiation by the transcription factor (TF) IIIB triggers restructuring of the DNA-bound Pol III. This restructuring can be mutagenic by itself and create mutational hotspots upstream of RNU genes.

Immunoglobulin kappa genes also exhibit long stretches of extreme hypermutability. In contrast to Pol III transcripts, however, sequencing quality metrics raise concerns about the reliability of SNVs in these genes (Supplementary Figure 11).

Transcription factor binding occurs at short scales and has been shown to be highly mutagenic in yeast and human cancers either because of blocked resection of ribonucleotide primers introduced by polymerase alpha, interference with the access of nucleotide excision repair, or altered DNA conformation^27,49–51^. First, we attributed TFBS activity to specific tissues by overlapping ChIP-seq signals with regions of open chromatin measured by DNase I hypersensitivity. In the majority of TFBS, Roulette predicts mutation rates accurately, confirming that the observed mutation rate elevation within TFBS is due to sequence context and regional features^52^ included in the Roulette model (Figure 5a). TFBS active in testis are a notable exception characterized by increases in the germline mutation rate over the background for most mutation types (Supplementary Figure 15a, b), with the strongest effect for T>G mutations (median increase across TFs is 1.59-fold, Figure 5a). This observation suggests a direct mutagenic effect of transcription factor binding.Interestingly, binding of SNPC4, the factor responsible for RNU transcription, has the strongest (6-fold) impact on mutation rate.

**Figure 5.**
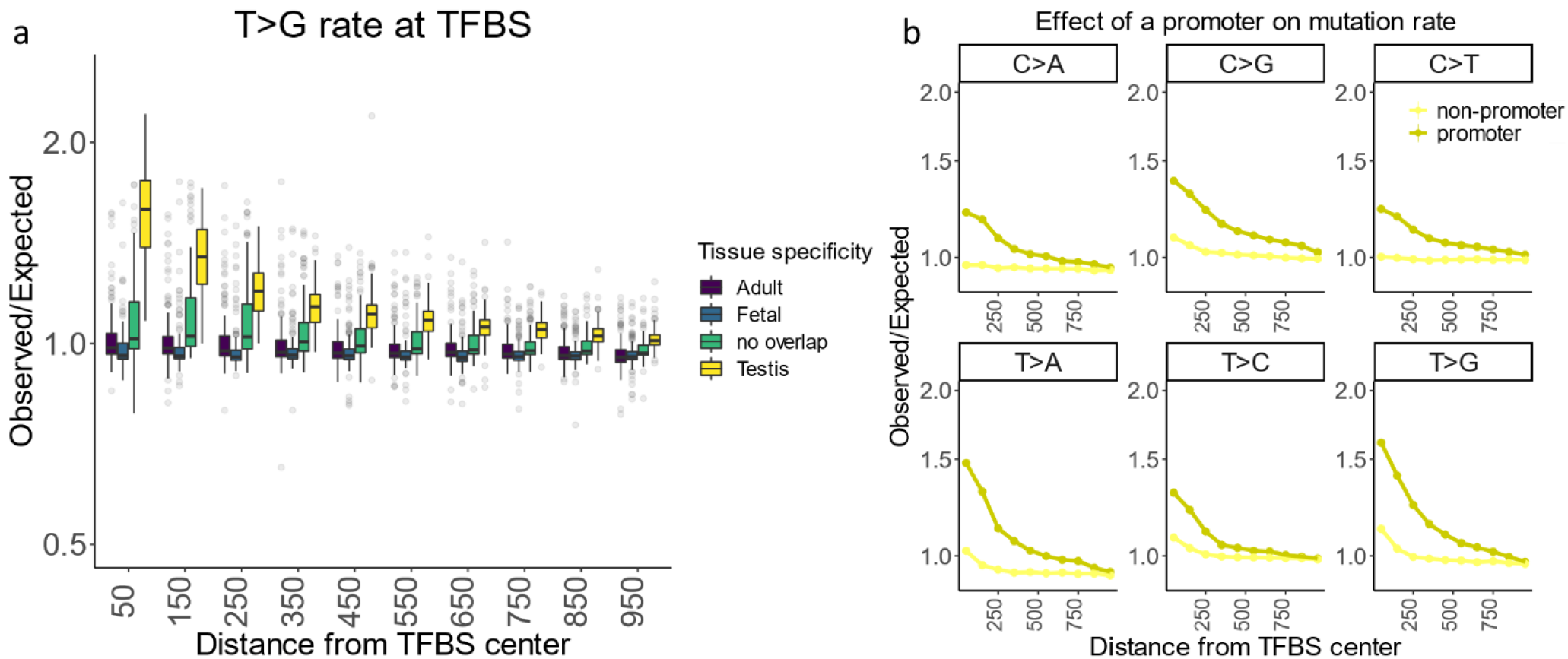
TFBS are prone to high mutation rate. a) Box plot for the observed to expected rate of rare T>G mutations across different transcription factors. Positions occupied with TF were annotated with chip-seq data. Tissues where TFBSs are active were determined through overlap with tissue specific DHS peaks. b) mutagenic effect of TFBS active in testis overlapping promoter (−2 kb upstream of transcription start site, dark yellow) or not (light yellow). c) and d) strand resolved observed to expected mutation rates at 100 nucleotide windows around TFBS centers in promoters.

Furthermore, we found that the higher mutation rates are almost exclusively restricted to testis-active TFBS in promoters (Figure 5b), and that TFBS overlapping multiple promoters have higher mutation rates than TFBS overlapping a single promoter (Supplementary Figure 16). To allow the application of Roulette to these genomic regions, we also provide mutation rates corrected for this TFBS effect (see methods, Supplementary Figure 17). Interestingly, similarly to germline mutations, UV-induced mutations at TFBS in melanoma also have different rates in and outside of promoters (Supplementary Figure 18)^51^.

As shown above, Roulette offers a significantly more accurate human mutational model and has demonstrated utility across different biological fields. Mutation rate estimates from the three analyzed models are made available here: http://genetics.bwh.harvard.edu/downloads/Vova/Roulette/. Future work may explain the sources of the demonstrated residual mutation rate variation, some of which may derive from evolving rates through time and variability between populations^53–55^.

## Code availability

Code used to perform the analysis is available at https://github.com/vseplyarskiy/Roulette.

## Data availability

Polymorphism data used in the study is freely available at https://gnomad.broadinstitute.org/. De novo mutations have been aggregated from supplementary materials to the refs 18 and 30.

Mutation rate estimates for autosomes http://genetics.bwh.harvard.edu/downloads/Vova/Roulette/. Shet values re-calculated with the help of Roulette could be found here http://genetics.bwh.harvard.edu/genescores/selection.html.

## Acknowledgements

We thank J. Wakeley and L. Fan for helpful suggestions on population genetics theory. We thank D.J. Balick for providing a forward Wright-Fisher simulator. This research was supported by National Institutes of Health grants R35-GM127131, R01-MH101244, U01-HG012009, R01-HG010372, and R01-HG010372 along with funding from NGM Biopharmaceuticals. D.J.L. was supported by NLM T15LM007092.

