## Supplementary materials for "A mutation rate model at the basepair resolution identifies the mutagenic effect of Polymerase III transcription"

**Methods**

*Genome sequencing data*

Roulette was fitted using whole-genome sequencing data from gnomAD v.3, representing 71,702 unrelated individuals^1^. For model validation, we used gnomAD v.2.1.1, a set of jointly called whole-exome sequencing data from 125,748 unrelated individuals with no known severe disorders (GRCh38 liftover version downloaded from <https://gnomad.broadinstitute.org/downloads>), UKBiobank whole genome-sequencing of 200,000 individuals, and *de novo* sequencing of 41,816 offsprings from Satterstrom *et al.*^2^, Halldorson *et. al.*,^3^ and An *et. al.*^4^. We also used gnomAD v.2.1.1 for population genetics analyses. Canonical transcripts from Ensembl v.104 were used to annotate transcription start sites, intron boundaries and all potential single nucleotide mutations as synonymous, missense, stop gained, splice donor, or splice acceptor variants. Observed variants were filtered to those annotated as PASS quality. For variants monomorphic in gnomAD, the number of successfully genotyped chromosomes (allele number) was not reported. We interpolate the allele number at monomorphic sites by using the average allele number among observed variants in the gene. *De novo* mutations were lifted over from GRCh37 to GRCh38.

*Implementation of the mutational model*

*1. Variables*

The Roulette mutational model was fitted using a set of known correlates of mutation rate in the human genome. For each non-coding autosomal position in the genome, we assessed the pentanucleotide context plus four additional adjacent nucleotides to the left and to the right of pentamer, the direction of replication, and the methylation level at CpG sites. To account for local mutation rate variation, we also included the estimated mutation rate in the 50 kb window of the site, conditional on mutation type and trinucleotide context. This accounts for context specific variation along the genome from sources not directly modeled. For sites within a gene body and in promoters, we run the model separately for reverse complementary mutations and to account for the potentially different mutational patterns we split the data into for equal-size bins by expression in testis.

We used the average effect of the surrounding nucleotides on mutation rate as an input variable to have a quantitative instead of categorical variable. For example, to predict a mutation in the underscored position with the following context A_-6_A_-5_C_-4_T_-3_ TG C>T GA T_3_C_4_C_5_A_6_ we used the observed value of μ(TG C>T GA |A_-6_)/ μ(TG C>T GA) as covariate for position. This ratio was separately calculated for each pentamer×compartment model, estimates for the modulating effects of surrounding nucleotides provided here (http://genetics.bwh.harvard.edu/downloads/Vova/Roulette/covariates/). All positions from -3 to -6 and 3 to 6 were treated similarly. Mutations within transcribed regions, intergenic regions, and promoters were considered separately because the effect of surrounding nucleotides differs between genomic compartments (Supplementary Figure 18).

The per-site direction of replication was obtained from ref.^5^ and averaged on a 10 kb scale. We aggregated fork direction into five equal size bins. For sites where the direction of the replication fork was not available, we assumed an absence of replication fork bias.

CpG methylation level for testis measured with bisulfite sequencing was obtained from ENCODE (<https://www.encodeproject.org/search/?type=Experiment&control_type!=*&status=released&perturbed=false&assay_title=WGBS&replicates.library.biosample.donor.organism.scientific_name=Homo+sapiens&biosample_ontology.term_name=testis>). We stratified CpG sites by a fraction of methylated reads into five methylation bins with incremental change of methylation by 0.2 and ran the model separately for each bin, because we are expecting different context effects for different methylation bins.

Mean expression level in testis was obtained from GTEx 8^6^.

As an additional variable to account for local variation in mutation rate, the average mutation rate in each trinucleotide context and each mutation type was calculated for sliding 50 kb windows with 10 kb steps. First, we calculated the genome-wide fraction of segregating mutations with trinucleotides using rare SNVs from gnomAD v3. To achieve independence between the local mutation rate and the site in consideration, we excluded the focal site from assessments of local mutation rate. Then, we normalized the fraction of segregated sites in each region against the genome-wide average. When an insufficient number of a specific trinucleotide was present in a region (< 10), the genome average mutation rate for this trinucleotide was assumed (covariate set to 1).

*2. Running the model*

All variables described above were used to predict the probability that a specific mutation was segregating as a rare SNV (< 0.1% frequency in gnomAD v3). Common variation was excluded because it is more sensitive to direct selection, background selection, and biased gene conversion. The model was trained using only non-coding sequences and low-quality sites were also omitted (see Methods section on filtering).

First we grouped sites into categories that we separately train our model, grouping accounts for 3072 pentamer-mutation pairs in each of three genomic compartments (intergenic regions, intronic regions, and promoters). We further split CpG sites into 5 bins by methylation and also separated promoter regions and gene bodies into five bins by expression level in testis. We then fit three separate models on a randomly selected 50% of sites within each of the resulting groups of sites. These were a logistic regression model with pairwise interactions, a logistic model with no interactions, and a model where the effects of sequence context and local mutability were estimated independently then multiplied. We compared log-likelihoods for all three models using the remaining 50% of sites unused in training and selected the model with the highest log-likelihood.

Logistic regression with pairwise interactions was applied with the following R-command:

*glm(*$SNVˇ$*~(N_-6_+ N_-5_+ N_-4_+ N_-3_+ N_3_+ N_4_+ N_5_+N_6_+Rep_dir+local_mutation_rate)^2, family = "binomial")*

Logistic regression without interactions was applied with the following R-command:

*glm(*$SNVˇ$*~(N_-6_+ N_-5_+ N_-4_+ N_-3_+ N_3_+ N_4_+ N_5_+ N_6_+Rep_dir), family = "binomial")*

$SNVˇ$ is the probability of not observing a segregating SNV site (see Methods section *“Poisson Transformation for Recurrence Correction”*), *N_x_* is the effect of the surrounding nucleotide **N** in position **x** relative to the focal site, *Rep_dir* is direction of replication, *local_mutation_rate* is the normalized rate of the same mutation in the overall trinucleotide context in the 50 KB window harboring a site. Note that sites that were used to train a model are in the same transcription level bin if they reside within gene bodies*,* and have similar methylation level if these sites are CpGs. The model fit by glm’s binomial family option is

$log(\frac{SNVˇ}{1-SNVˇ})=X\beta+\varepsilon$,

where $X$ is a matrix of all observed covariates, including interactions, and $\beta$ is the estimated effect of each on SNV occurence.

A simpler log-link model was included for pentamer-compartment categories with few observed SNVs. For instance, certain sequence contexts that are rare in promoters. For this model we estimated the effect of each covariate independently (no multiple regression), and multiplied the odds ratios to obtain the probability of not observing an SNV:

$SNVˇ$*~N_-6_* N_-5_*N_-4_*N_-3_* N_3_* N_4_* N_5_* N_6_* local_mutation_rate*

The likelihood of this model was then also compared to the case of a constant mutation rate (one parameter) within that category of sites. The fraction of the genome fit best by each model in the held-out 50% of sites is shown in Supplementary Figure 21.

*3. Poisson Transformation for Recurrence Correction*

Due to recurrent mutations at high mutation rate sites in a large sample like gnomAD, there is not a linear relationship between the SNV probabilities fit using logistic regression and the underlying mutation rates. We develop and justify a simple Poisson transformation to recover estimates on the mutation rate scale. This approach applies the classic infinite-sites model to individual sites in the genome. A detailed mathematical justification for this approach can be found in Wakeley et al. 2023.

Assuming constant mutation rate $\mu$ along a particular lineage of length $t$, the infinite-sites model states that the number of mutations in that lineage will be $Pois\left( \mu t \right)$. Let $S$ be the number of mutations in the genealogy of a sample of individuals. It follows that $S\sim Pois\left( \mu T_{tot} \right)$, where $T_{tot}$ is the total length of genealogy, because $T_{tot}$ is the sum over all branches in the genealogy. However, $T_{tot}$ varies along the genome due to the inherent stochasticity of the coalescent process, and is also affected by linkage to selected sites. For a non-recombining locus, a category applicable to single sites in the human genome, the variance of S is $Var\left( S \right)=\mu E\left( T_{tot} \right)+\mu^{2}Var\left( T_{tot} \right)$ (Watterson 1975)^7^. The Poisson approximation is therefore applicable when $\mu^{2}Var\left( T_{tot} \right)$ is small. In a large sample from a constant-size population $Var\left( S \right)=\theta\frac{\sum_{i=1}^{n-1} 1}{i}+\pi^{2}\frac{\theta^{2}}{6}$, where $\theta=4N\mu$, $N$ is the effective population size, and $n$ is the sample size. Since $\theta\ll1$, for large $n$, $Var\left( S \right)$ approaches $E\left( S \right)$ and the distribution of $S$ becomes approximately Poisson (Ewens 2004, pg. 299)^8^. Indeed, even the distributions of the number of low counts SNVs (singletons, doubletons, etc.) are approximately independent and Poisson distributed as the sample size becomes very large^9^. The variance in $T_{tot}$ is therefore negligible.

The recent growth of the human population results in genealogies with a greater proportion of $T_{tot}$ residing in branches with fewer descendants compared to constant size populations. Summing over a larger number of branches with fewer descendants decreases $Var\left( T_{tot} \right)$ compared to a constant-sized population, increasing the accuracy of the Poisson approximation. Therefore, for a sample as large as gnomAD v3 (71,702 individuals), we can expect $SPois\left( \mu T_{tot} \right)$. Under the assumption that S is Poisson, we have $P_{0}\left( \mu\right)=exp\left( -T_{tot}\mu\right)$, where $P_{0}$ is the probability of a site experiencing no mutation in the history of the sample. This is equivalent to a site being monomorphic as long as back mutations to the ancestral state are sufficiently rare. This condition is satisfied because the occurrence of any nested rare mutations at realistic human mutation rates is negligible^10^. Therefore, $\mu=-log\frac{\left( P_{0}' \right)}{T_{tot}}$, yielding a linear relationship between $log\left( P_{0} \right)$ and $\mu$. This provides a scaling between the monomorphic probabilities estimated by logistic regression models and the underlying mutation rates.

To apply mutational models developed by other groups on the gnomAD dataset, which is prone to recurrent mutations, we needed to transform them to the $P_{0}$ scale. To do so, we estimated a scaling parameter *T_tot_* by first binning continuous mutation rate estimates and then solving the following equation:

$\sum_{i} L_{i}*e^{{-T_{tot}\mu}_{i}}=L-P$

Where *i* indexes the mutation rate bin, $L_{i}$ is the number of sites in mutation rate bin *i*, $\mu_{i}$is the mean mutation rate predicted by the model within bin *i*, L is the overall number of sites and P is the overall number of polymorphic sites.

After finding $T_{tot}$, we calculate the expected number of polymorphic sites by calculating $\sum_{i} L_{i}\left( 1-e^{{-T_{tot}\mu}_{i}} \right)$.

*4. Filtering*

We marked regions and individual sites as low quality based on quality criteria from gnomAD, abnormal density of SNV sites, and on suspicious patterns of recurrence.

More specifically, we classified sites as low quality if Umap100 mappability was below 0.5, if the site overlapped with a long (>50 nucleotide) simple repeat, if in a hundred nucleotide window the mean ReadPosRankSum was above 1, or if the number of segregating SNVs in a hundred nucleotide window was zero.

After we obtained predicted mutation rates for every site, we noticed that within some rate categories the site frequency spectra (SFS) has an abnormally high fraction of high frequency variants. We noticed that this problem is context specific. To separate problematic sites for each pair of pentamer and mutation rate we calculated fraction of high frequency SNVs [MAF >0.005 and MAF <=0.2] and compared it to the average fraction of high frequency SNVs across all mutation types; the pentamer was labeled unreliable if the fraction of high frequency sites is 1.5-fold higher than the mutation rate-controlled average (See Supplementary Figure 19). Most sites masked this way belong to repetitive contexts like AAATT>T, TTAAA>T, AATAT>A or CTCTA>A (Table S3).

*5.Validation*

I. Pseudo R^2^

We want to measure how well estimated mutation rates fit *de novo* mutation data. Ideally, this measure should allow us to make a fair comparison between different mutation rate models, should be insensitive to sample size, and should give some sense of distance from a theoretical optimum. We followed Nagelkerke (1991) who defined $R^{2}$ for general regression models. One possible definition is $R^{2}=1-exp\left( \frac{-2}{n}\left[ l\left( \hat{\beta} \right)-l\left( 0 \right) \right] \right)$, where $l\left( \hat{\beta} \right)$is the log-likelihood of the model and $l\left( 0 \right)$ is the likelihood under some null. For discrete outcomes the maximum value of this $R^{2}$ is $max\left( R^{2} \right)=1-exp\left( \frac{2}{n}l\left( 0 \right) \right)$.

Nagelkerke (1991) proposed ${R´}^{2}=\frac{R^{2}}{max(R^{2})}$ as a better metric for the proportion of explained variation. However, the mutation process is inherently stochastic. The $max\left( R^{2} \right)$ above assumes that all observations could be predicted perfectly, whereas perfect knowledge of mutations rates would never do this. The true mutation rate model would assign a probability $\mu_{i}$ to each potential mutation. If the log-likelihood of the data under this model is $l\left( \beta\right)$, the we can define the maximum $R^{2}$ as

$max\left( R^{2} \right)=1-exp\left( \frac{-2}{n}\left[ l\left( \beta\right)-l\left( 0 \right) \right] \right)$.

$\frac{1}{n}l\left( \beta\right)=\frac{1}{n}log\sum\left( \mu_{i}^{x_{i}}\left( 1-\mu_{i} \right)^{1-x_{i}} \right),$

where the sum is across sites and $x_{i}$ indicates whether a mutation is observed or not. It is possible that $R^{2}$ exceeds the theoretically optimal $max\left( R^{2} \right)$ by chance when the sample size is small, so empirical values of ${R´}^{2}$ greater than one are possible.

Of course, we do not know the true $\mu_{i}$. However, we can use knowledge about the overall distribution of mutation rates. If we know $f\left( \mu\right)$,

$E\left[ \frac{1}{n}l\left( \beta\right) \right]\to\int\left( \mu log\mu+\left( 1-\mu\right)log\left( 1-\mu\right) \right)f\left( \mu\right)d\mu$,

for large n. The appropriate null model is $\underline{\mu}$, the genome-wide average mutation rate. The maximum R^2^ would then be approximately

$max\left( R^{2} \right)=1-exp\left( -2\left[ \int\left( \mu log\mu+\left( 1-\mu\right)log\left( 1-\mu\right) \right)f\left( \mu\right)d\mu-\underline{\mu}log\underline{\mu}-\left( 1-\underline{\mu} \right)log\left( 1-\underline{\mu} \right) \right] \right)$.

One implication here is that we cannot say how far we are from the optimal mutation rate model without making some assumption about the overall distribution of mutation rates $f\left( \mu\right)$. This is because it is not possible to distinguish model errors from mutational stochasticity. As $f\left( \mu\right)$, we use the distribution of rates fit by the Roulette model.

We calculate pseudo R2 as ${R´}^{2}=\frac{R^{2}}{max}\left( R^{2} \right)$ where

$R^{2}=1-exp\left( \frac{-2}{n}\left[ \sum log\hat{\mu_{i}}{}^{x_{i}}\left( 1-\hat{\mu_{i}} \right)^{1-x_{i}}-log\underline{\mu}{}^{x_{i}}\left( 1-\underline{\mu} \right)^{1-x_{i}} \right] \right)$

and $max\left( R^{2} \right)$ is as defined above.

We can also apply pseudo R^2^ to gnomAD v2.1.1 polymorphism data by replacing $x_{i}$ with $y_{i}$, which represents whether a site is polymorphic or not. We replace $\mu_{i}$ with $p\left( \mu_{i} \right)=\left( 1-e^{{-k\mu}_{i}} \right)$ (from the section on Poisson Transformation for Recurrence Correction), where ${p(\mu)}$ is the probability of a site being polymorphic given mutation rate $\mu$. Therefore, we use:

$max\left( R^{2} \right)=1-exp\left( -2\left[ \int\left( p\left( \mu\right)logp\left( \mu\right)+\left( 1-p\left( \mu\right) \right)log\left( 1-p\left( \mu\right) \right) \right)f\left( \mu\right)d\mu-(p\left( \underline{\mu} \right)logp\left( \underline{\mu} \right)-\left( 1-p\left( \underline{\mu} \right) \right)log\left( 1-p\left( \underline{\mu} \right) \right) \right] \right)$

For bootstrapping, we sample synonymous variants with replacement 1,000 times. Figure 2a represents the 95% confidence interval of pseudo-R^2^ over the samples. For figure 2b, we subtract pseudo-R^2^ of Roulette from that of other models for each sample. Figure 2b shows the 95% confidence interval of this pseudo-R^2^ difference.

II. *Residual variance*

In addition to the pseudo-R^2^ analysis described above, we developed an alternative method to estimate the proportion of mutation rate variation not explained by a given model. This method separates sites by whether that SNV was observed in gnomAD v3 (test set). Within each mutation rate bin defined for that model we compared the numbers of *de novo* mutations observed in an independent data set between monomorphic and polymorphic sites. If mutation rates are estimated perfectly, there will be little to no difference in the proportion of sites with *de novo* mutations, whereas large differences would indicate substantial residual variance in true mutation rates.

To use stratification by SNV status to estimate residual mutation rate variances, we assume that the distribution of mutation rates within bin $i$ is $f_{i}\left( \mu\right)$. The distribution of rates conditional on observing a monomorphic site is

$f_{i}\left( \mu|noSNV \right)=\frac{p\left( no SNV|\mu\right)f_{i}\left( \mu\right)}{p\left( no SNV \right)}$. The probability of observing no SNV is approximately $e^{-\mu T_{tot}}$ (see Methods, *Poisson Transformation for Recurrence Correction*), and the mutation rate in *de novo* data will be equal to the average rate for this group of sites:

$E_{i}\left( \mu|no SNV \right)=\int\mu f_{i}\left( \mu|no SNV \right)d\mu=\frac{\int\mu e^{-\mu T_{tot}}f_{i}\left( \mu\right)d\mu}{\int e^{-\mu T_{tot}}f_{i}\left( \mu\right)d\mu}$.

The equivalent expression for sites with no SNV is

$E_{i}\left( \mu|SNV \right)=\int\mu f_{i}\left( \mu|SNV \right)d\mu=\frac{\int\mu(1-e^{-\mu T_{tot}}{)f}_{i}\left( \mu\right)d\mu}{\int(1-e^{-\mu T_{tot}})f_{i}\left( \mu\right)d\mu}$

After doing the same for polymorphic sites we can calculate the ratio for the excess of *de novo* mutations expected at polymorphic sites:

$\rho_{i}=\frac{E_{i}\left( \mu|SNV \right)}{E_{i}\left( \mu|noSNV \right)}=\frac{E_{i}\left( \mu\right)-E_{i}\left( \mu e^{-\mu T_{tot}} \right)}{E_{i}\left( \mu e^{-\mu T_{tot}} \right)}\frac{E_{i}\left( e^{-\mu T_{tot}} \right)}{1-E_{i}\left( e^{-\mu T_{tot}} \right)}$.

This is because the expected mutation rate in a bin is proportional to the probability that a *de novo* mutation is observed.

Similarly to the pseudo-R^2^ analysis, it is necessary to make some assumption about the distribution of mutation rates, this time the residual distribution within each bin. We assumed a log-normal distribution of mutation rates and re-estimated the mean within each bin using the total de novo mutation count. $\rho_{i}$ thus depends on two parameters, the log-scale standard deviation of mutation rates $\sigma_{i}$ and $T_{tot}$. These two parameters are identifiable within each bin given the observed *de novo* ratio and proportion of polymorphic position. In theory $T_{tot}$ should be the same for each bin as all share a common population history. In practice we estimate $T_{tot}$ separately for each bin and confirm that it does not vary too widely (Supplementary Figure 23).

We fit $\sigma_{i}$ and $T_{tot}$for each bin using a grid search to identify the point where

$\left| logE_{i}\left( P_{1}|\sigma_{i},T_{tot} \right)-log\hat{P_{1}} \right|+\left| logE_{i}\left( \rho|\sigma_{i},T_{tot} \right)-log\hat{\rho_{i}} \right|$

is minimized, where $\hat{P_{1}}$ is the observed probability a site is polymorphic in that bin and $\hat{\rho_{i}}$ is the observed *de novo* ratio. Expected proportions of polymorphic sites and *de novo* ratios were calculated by sampling large numbers of mutation rates from each possible log-normal distribution in the grid. Estimates of $\sigma_{i}$ fell within the range [0, 1.5] and were transformed to a linear scale using

$\hat{\sigma_{\mu,i}}{}^{2}=\left( e^{\sigma_{i}^{}^{2}}-1 \right)\hat{\mu_{i}}{}^{2}$.

The residual variance contributed by each bin was then computed as

$V_{i}=\frac{p_{i}\hat{\sigma{}_{\mu,i}}{}^{2}}{\sum p_{i}\left( \hat{\sigma{}_{\mu,i}}{}^{2}+\left( \underline{\mu}-\hat{\mu_{i}} \right)^{2} \right)}$.

III. Per-Gene Z-score

To calculate the Z-score, we assume that the number of polymorphic sites in a gene follows a Poisson-binomial distribution. The expected number of polymorphic sites is $\sum_{i\in L} P_{1}\left( \mu_{i} \right)$. The variance is the sum $\sum_{i\in L} \left( 1-P_{1}\left( \mu_{i} \right) \right)P_{1}\left( \mu_{i} \right)$ over set of sites $L$, where $P_{1}\left( \mu_{i} \right)=1-e^{{-k\mu}_{i}}$.

*6. Corrections to the model and application to the X chromosome*

Comparing the predicted and observed number of rare SNVs shows that some regions have much higher mutation densities than expected. While we attributed some of these mutational hotspots to transcription by polymerase III, for other regions the biological etiology was unclear. To recalculate the mutation rate in hypermutable regions we ran additional logistic regressions among hotpots – 100 nucleotide windows with more than 75 rare SNVs. For this regression we used previously estimated mutability and mutation type to predict an adjusted hypermutable mutation rate. We applied a similar procedure to adjust for higher mutation rates at transcription factor binding sites (TFBS), whereas as with additional variables we used the type of transcription factor, distance from the center of the CHIP-seq peak, overlap with a promoter and tissues where the factor is active. We provide both adjusted and initial mutation rate predictions. We used an analogous procedure to re-calibrate mutation rate in contexts that have atypical SFS.

*7. Genomic features*

Chip-seq tracks were downloaded from the Vorontsov *et. al.*^11^ and only category A data (highest quality) were used. Chip-seq signal from different overlapping TFBS were counted independently (Figure 4).

DHS tracks were downloaded from ENCODE (https://www.encodeproject.org/search/?type=Experiment&assay_title=DNase-seq). We aggregated DHS peaks from 4 adult tissues (lung, stomach, leg muscle, brain) to obtain the “adult” DHS track and from 4 embryonic/fetal tissues (fibroblast, placenta, large intestine, stomach) to obtain the “fetal” DHS track, finally we aggregated two tracks from testis to obtain the testis track. In order to obtain peaks private to “fetal” and “adult” tissues, we excluded peaks that overlapped between them or with “testis”. Testis, in contrast, includes ubiquitous peaks, but using the private testis track does not change our results.

Annotations of active genes and pseudogenes that are transcribed by polymerase III (tRNA, snRNA, vault RNA, RNA component of 7SK nuclear ribonucleoprotein, ribonuclease P RNA component H1, ribosomal RNA, Ro60-associated Y RNA) were downloaded from HUGO Gene Nomenclature Committee (<https://www.genenames.org/data/gene-symbol-report/>). It has been reported that some ALU repeats are also transcribed by polymerase III^12^, and we downloaded coordinates of such ALU elements from ref. ^12^.

We used Ensembl for genic annotations and definition of promoters. We defined promoters as a region 0 to 2 KB upstream of the CDS containing the gene. We do not exclude regions that match our definition of promoters for more than one transcript.

*8. Demographic Inference*

In order to determine whether mutation rate estimates from Roulette are sufficient to capture distortions to the site frequency spectrum due to recurrent mutation, we fitted a demographic model and assessed how well it matched the observed SFS in each mutation rate bin. We based our demographic model on those presented by Gao and Keinan (2016)^13^ and Gazave *et. al.,* (2014)^14^. The Gazave model starts with a constant population size, followed by two bottlenecks (one of which is the out-of-Africa bottleneck), and ends with a recent population growth. Gao and Keinan (2016) used the parameters from Gazave et al. (2014), but re-estimates the parameters for the recent exponential growth phase. They conclude that the best demographic model has a faster-than-exponential growth, with growth speed parameter $b=1.12$ ( $b=1$ is equivalent to exponential growth). We keep the model from Gao and Keinan (2016), but refit the parameters $b$ and the initial growth rate $g$, which they estimated as 0.0055.

We use a Wright-Fisher simulator used in Weghorn *et. al.,* (2019)^15^ to simulate the site-frequency spectrum a dense grid of $b=\left[ 1.1,1.3 \right]$ and $g=\left[ 0.005,0.006 \right]$. Though this may seem like a narrow range, the final population size varies from 0.7 million to 40 million. Here, the log-likelihood is computed as:

$\left( b,g \right)=\sum_{i=0}^{56885} C_{i}*logE\left[ b,g \right],$

where $C_{i}$ is the folded synonymous allele counts for non-Finnish Europeans (NFE) in gnomAD v.2.1.1 whole exomes dataset and $E\left[ b,g \right]$ is the expected folded SFS given parameters *b* and *g*. Under this framework we find  $\left( b^,g^ \right)$ that maximizes the above likelihood.

We utilize the full mutation rate information to find the maximum log-likelihood. For each parameter, we simulate the SFS for all the mutation rate bins and calculate the log-likelihood for each mutation rate bin. Then, we sum up the log-likelihood over the bins and find the growth parameters that maximizes this summed likelihood.

Then, to compare how well the SFS fits within different possible mutation rate bins we re-fit μ using the maximum likelihood demographic parameters. We do this for all Roulette bins used in other analyses, as well as for low and high-rate defined as [1.3e-09, 3e-09] and [1e-07, 2.8e- 07]. The Wright-Fisher simulations allow for recurrent mutation, so the SFS changes shape as the mutation rate increases. We measure the fit to the shape of the SFS by calculating the likelihood conditional on sites being polymorphic by removing the zero bin and normalizing the remaining expected SFS. We evaluate the information added by Roulette’s fine-scale mutation rate estimates by comparing the conditional likelihoods of the low and high-rate fits to the μ fit specifically to that bin.

To evaluate the information added to demographic modeling by high rate sites we compute the average per-polymorphism contribution to the log-likelihood. We calculate the likelihood using the first 40 entries of the SFS under a model of pure exponential growth with recurrent mutation (Wakeley et al. 2022). As an example, we use the best fitting parameter $\alpha=N_{0}\frac{r}{n}\approx6$ to the rare SFS in the gnomAD v2 data, where $N_{0}$is the current population size, $r$ is the per-generation growth rate, and $n$ is the sample size. The parameter $\theta_{0}=4N_{0}\mu$ was matched to estimates in Roulette bins to provide comparison points at 3e-09, 3e-08, and 3e-07.

*9. Estimating fraction of ancestral variants in training dataset*

When fitting Roulette we assumed that alternative alleles with frequency < 0.001 were derived and therefore represented mutation events from the reference to the alternative state. This assumption can be violated if an alternative allele observed at frequency x is really an ancestral allele at frequency 1-x. In order to bound the fraction of rare variants (MAF < 0.001) used in training that are ancestral, we used the demographic fit of non-Finnish Europeans (NFE) mentioned above. We simulated the unfolded SFS of a high mutation rate class (3e-07 mutations per site per generation) using the Wright-Fisher simulator used for the demographic analysis. However, the gnomAD v3 dataset used to fit Roulette includes individuals with ancestry labels other than NFE, most of whom are of African descent. Since it was not feasible to fit another, more complex, demographic history, we used simulations of an equilibrium population to capture a greater range of potential SFS. The probability the minor allele is ancestral will be greatest for the highest frequency under consideration, so we report values for 0.001. For the high mutation rate class, we get that 1.25*10^-5^ and 3.09*10^-5^ of the rare variants are ancestral in NFEs and in equilibrium population, respectively.

*10. Validation of outlier gene classes*

Analyses of observed SNV counts and Roulette expectations in 100 nucleotide windows identified regions with potentially high mutability. We took a closer look at three large classes: IGK, RNU, and tRNA genes. RNU and tRNA are transcribed by Pol III, suggesting a shared mutagenic mechanism. We used population allele frequencies, *de novo* mutations, and quality scores to determine whether elevated SNV counts were due to true hypermutability.

Supplementary Note 1

1. Inferring mutation rates from the distribution of allele frequencies

The shape of the SFS for low-count alleles is strongly dependent on the mutation rate due to recurrent mutation (cite Harpak et al.). Roulette estimates are sufficiently accurate that it is reasonable for us to assume a single mutation rate within each defined bin when predicting the shape of the SFS for alleles as those sites (Figure 3) (cite Wakeley et al.). Given this, we can estimate the distribution of mutation rates in a group of sites by modeling its SFS as a mixture of the SFS shapes observed in each Roulette bin. To avoid issues with variant ascertainment, we only fit the distribution of frequencies conditional on an SNV being observed in the sample.

Let p(x) be the observed SFS at some sites of interest, e.g. those in RNU genes, and let p_μ_(x) be the SFS observed in a mutation rate bin with mean μ. p(x) can be modeled as a mixture of mutation rates: $p(x)=\sum\pi_{\mu}p_{\mu}(x)$, where $\pi_{\mu}$ is the proportion of *observed* variants with mutation rate μ and the quantity which we want to infer. In application to gene classes, we only used every fifth Roulette bin (21 total) to reduce overfitting. We estimated $\pi_{\mu}$ using maximum likelihood and the basinhopping algorithm with L-BFGS-G as the minimizer, as implemented in scipy. We also included a smoothing penalty on the multinomial transformation of $\pi_{\mu}$: $c\times\sum_{i=1}^{n_{bins}-1} {(\beta_{i}-\beta_{i+1})}^{2}$. We applied this model to the observed SFS in IGK, RNU, and tRNA genes with both no (c=0) and relatively strong (c=1000) smoothing. To bin the SFS, we used counts 1-5 without binning and variants were binned logarithmically with base 3 above that. Details of the analysis are provided at <https://github.com/vseplyarskiy/Roulette/blob/main/population_genetics/high_rate_gene_context_comparison.ipynb>.

Both the smoothed and unsmoothed analyses indicated a mix of intermediate and high mutation rate SNVs in RNU and tRNA genes (Supplemental Figure 14, Figure 4c,d). The SFS in IGK genes was less compatible with a high proportion of hypermutable sites.

2. *de novo* mutation rate conditional on SNV status

Using the same approach as for the residual variance, we separated sites based on their Roulette bin and whether they contained an SNV in the population sequencing data (gnomAD v3). We also separated transition and transversion mutations for tRNA and RNU genes. We then estimated de novo mutation rates for each group of sites. No *de novo* mutations were reported in IGK genes, consistent with the SFS-based prediction of little hypermutability. Exact Poisson confidence intervals were calculated using the R package “exactci”.

3. The distribution of quality metrics in hypermutable gene classes

To evaluate variants within hypermutable regions we used three quality metrics provided by gnomAD. Allelic balance is determined by the relative count of reads with the alternative and reference alleles. This should be concentrated around 50% for true germline variants, while variants representing somatic mutations will show up at frequencies <50% for the derived allele. We also examined mapping quality scores, which reflect the relative mapping likelihoods of alternative versus reference reads, in order to diagnose mis-mapping artifacts. These are shown in Supplementary Figure 22 and indicate that mapping quality is better in RNU and tRNA genes compared to the genomic background on chromosome 21, while scores in IGK are worse. Finally, we look at an overall allele-specific variant quality score (AS_VQSLOD) to capture any other factors.

**Supplementary Figures**

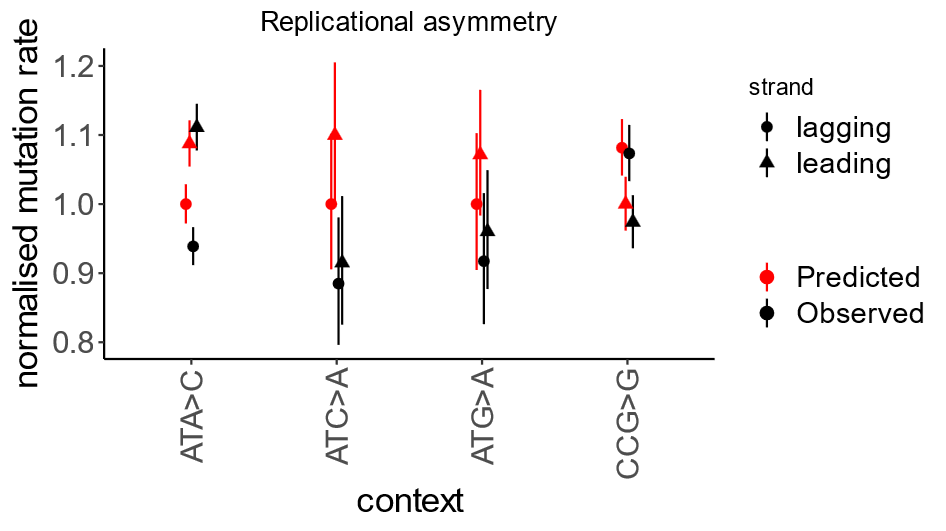

**Supplementary Figure 1. Effect of replication fork direction on the rate of rare synonymous SNVs.** Four contexts with strongest replication asymmetry. Mutation rate calculated for the regions with the strongest replication fork polarity (top quartile). Mutation rate is relative to the least mutable strand.

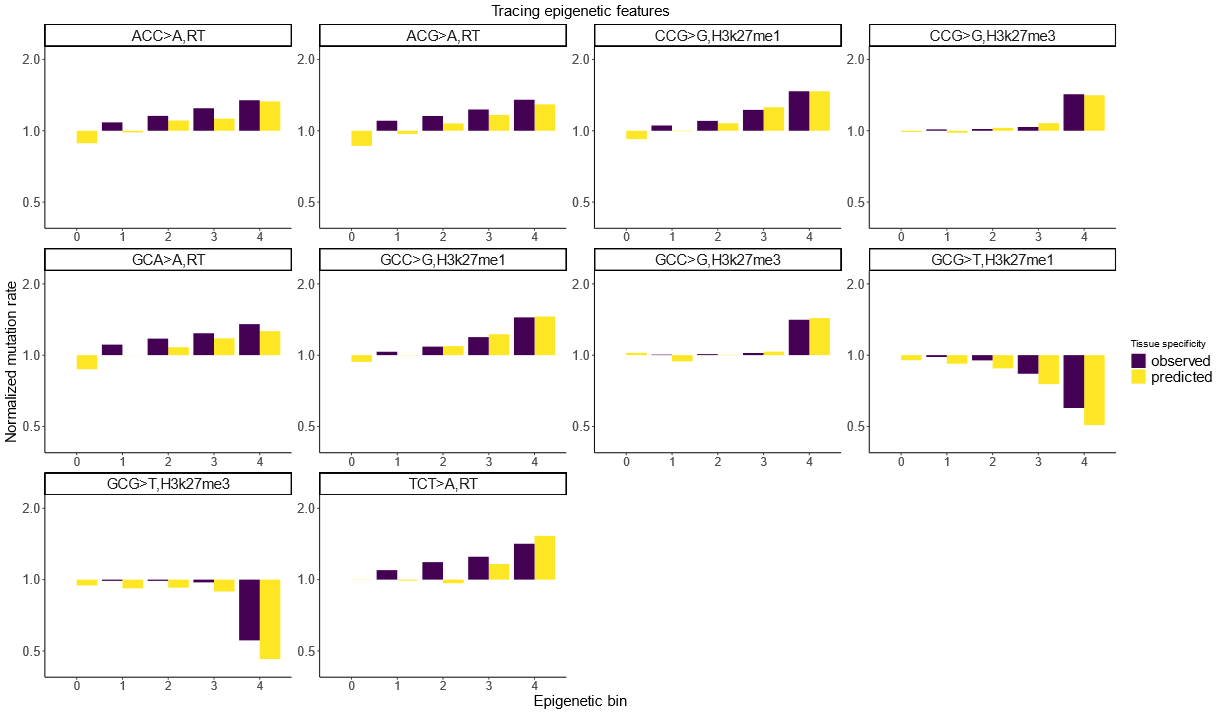

**Supplementary Figure 2. Roulette captures mutation rate variation associated with epigenetic features.** Ten pairs of mutation type and epigenetic features with the strongest effects on mutation rate. To generate bins, we subdivided the genome into five equal size bins by the value of genomic features and then calculated observed and expected mutation rates for each trinucleotide context among synonymous sites. This test was performed on synonymous SNVs and mutation rates were normalized to the rate observed in the first epigenetic bin. RT stands for replication timing. Overall, we analyzed the effect of replication timing, H3k27me3, H3k27me1, and recombination.

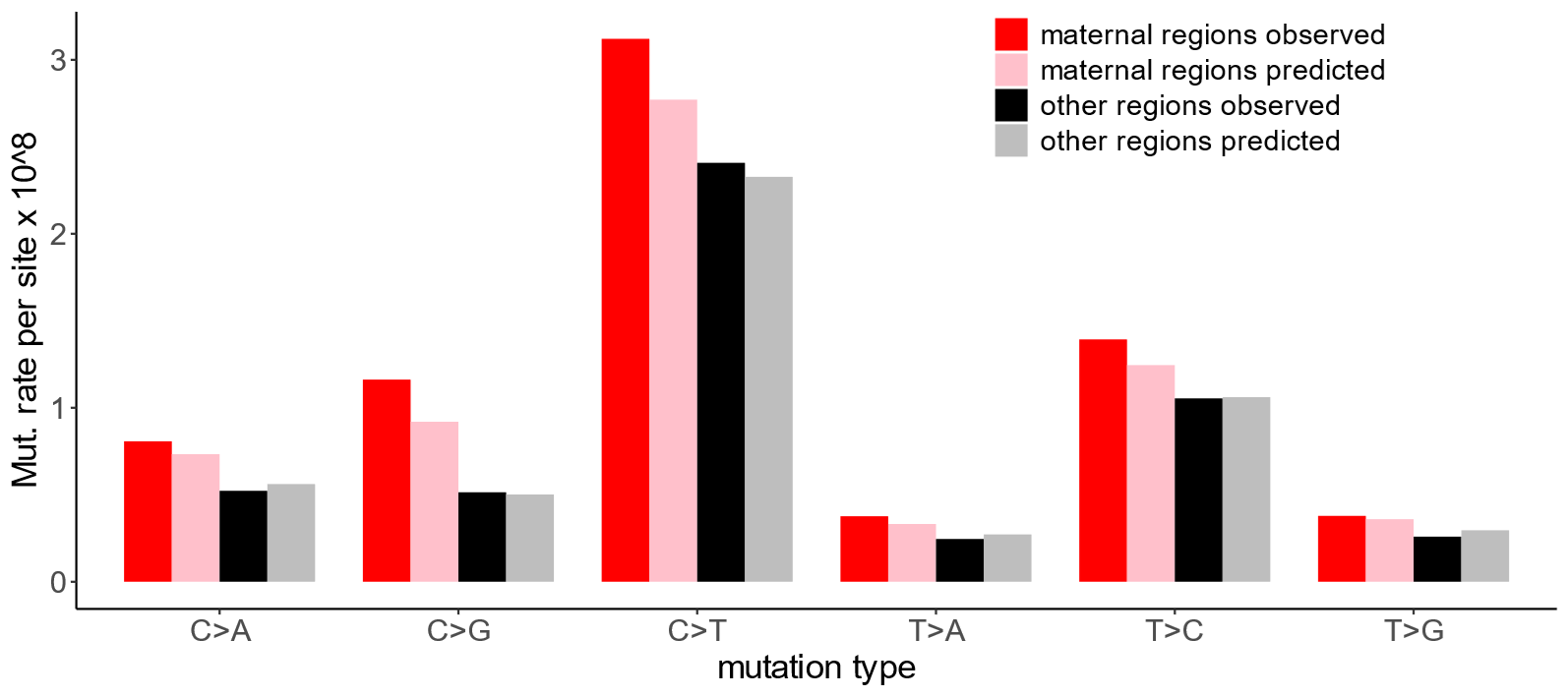

**Supplementary Figure 3. Roulette captures accelerated mutation rate in “maternal” regions.** *De novo* mutation rate inside and outside of maternal regions. Maternal regions are defined as in Seplyarskiy *et. al.,* 2021*.*

**Supplementary Figure 4. Roulette predicts the rate of triallelic SNVs.** Multiple derived alleles could co-occur in the same genomic site. Using Roulette, we predicted the probability of a site to contain two derived variants simultaneously (triallelic site) by multiplying the probabilities of each derived allele (this is the correct procedure if derived alleles accumulated independently). In contrast to early studies of multiallelic variants we do not find deviation from independence.
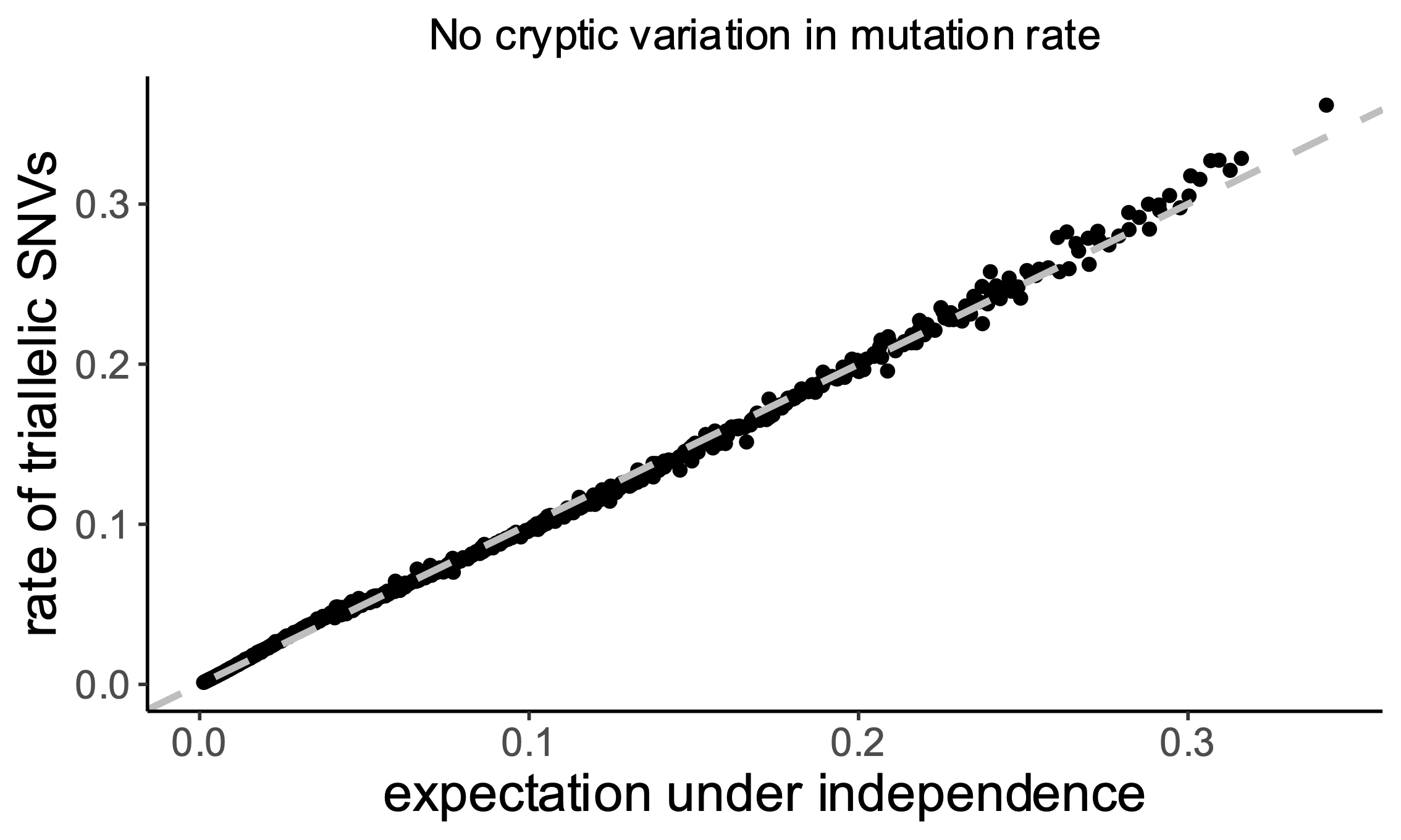

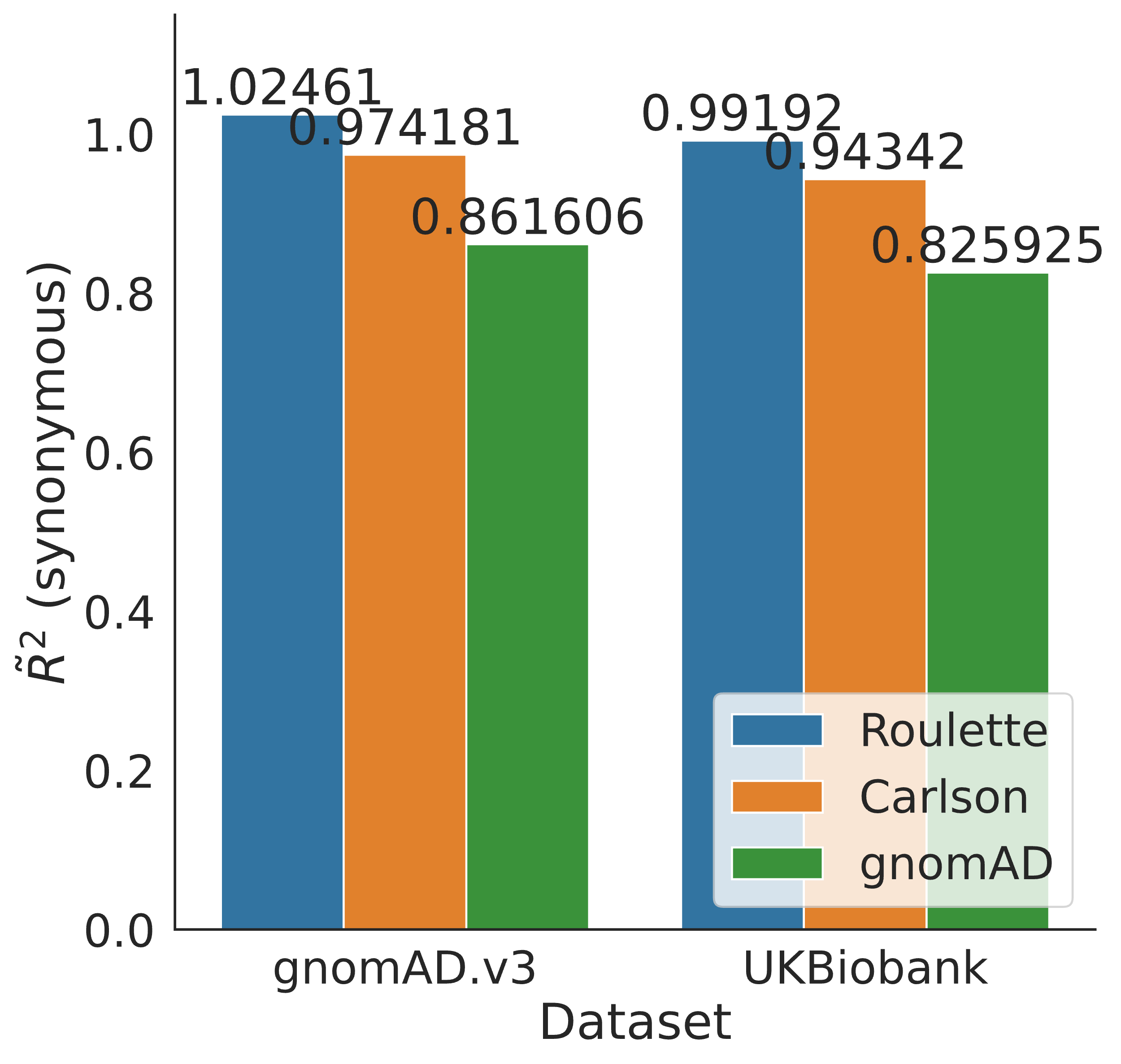

**Supplementary Figure 5. pseudo-R^2^ for noncoding regions.** pseudo-R^2^ is calculated for noncoding regions for two datasets: gnomAD.v3, and UK Biobank. Since Roulette was trained on noncoding variants from the gnomAD.v3, it is expected that Roulette performs better for noncoding variants than synonymous variants. *De novo* sequencing and UK Biobank population sequencing is an independent dataset from trained data.

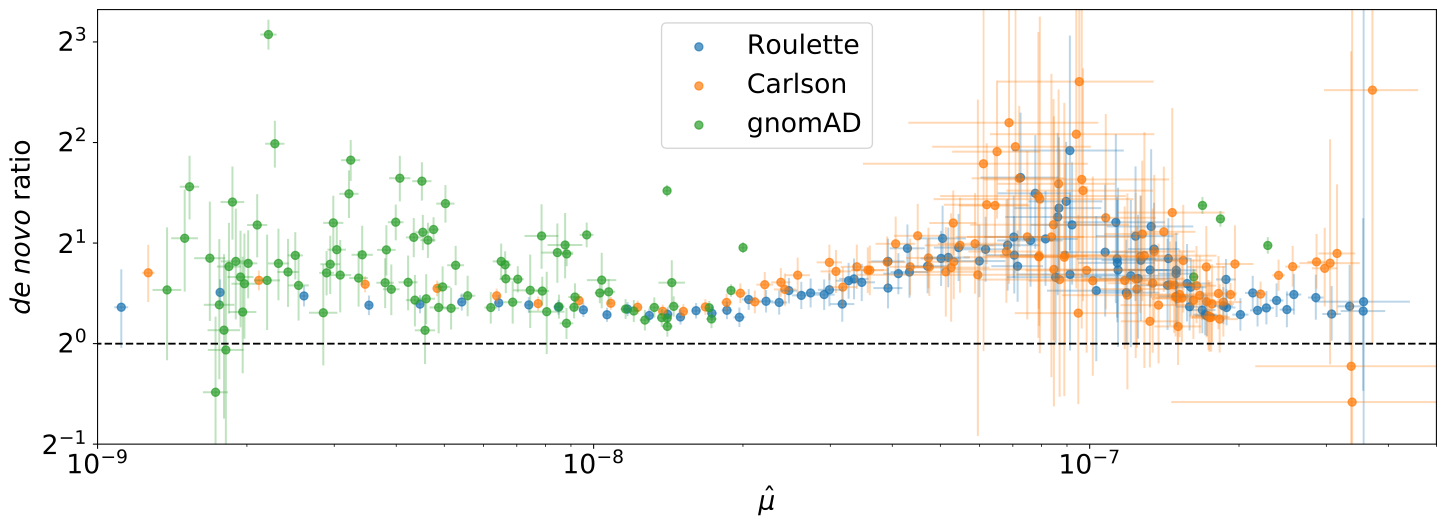
 **Supplementary Figure 6. A greater number of *de novo* mutations at sites with observed SNVs.** Sites were divided into mutation rate bins for the three different models. *de novo* mutation rates were calculated from whole-genome family sequencing data. Horizontal bars represent 95% Poisson confidence intervals for the *de novo* mutation rate within each bin. Vertical bars represent 95% confidence intervals for the ratio of Poisson rates between SNV and non-SNV sites within each bin.

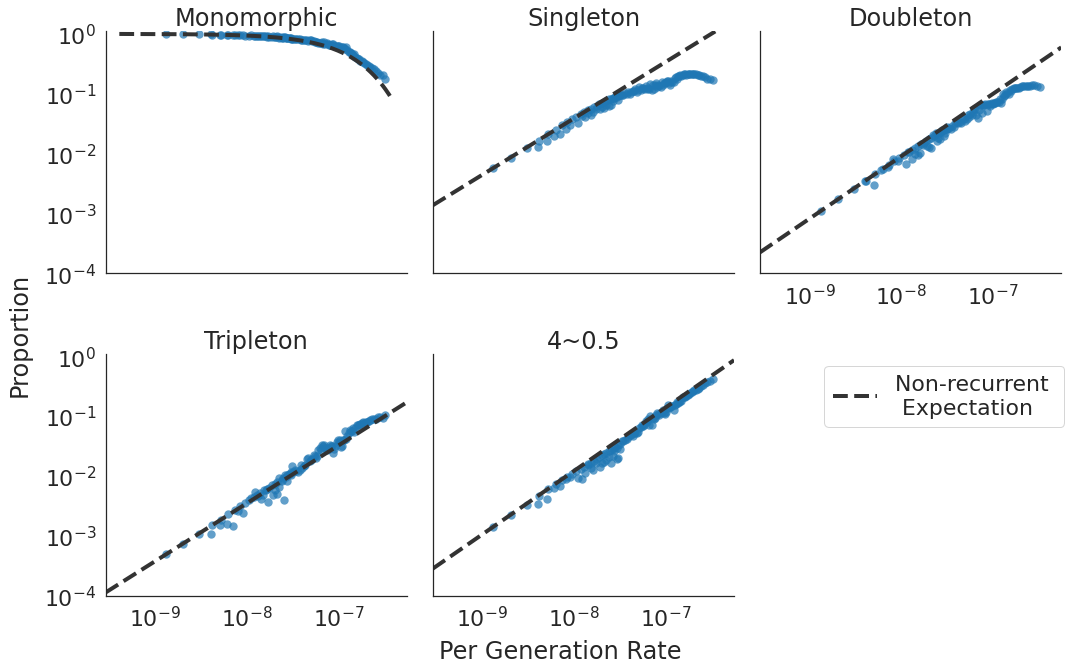

**Supplementary Figure 7. Recurrence breaks the expectation under non-recurrence.** Proportion of sites in five different classes: monomorphic sites, singletons, doubletons, tripletons, and other SNVs with higher allele counts. X-axis shows the per-generation mutation rate, as estimated by Roulette. The dotted line is the expected trend under the infinite sites model.

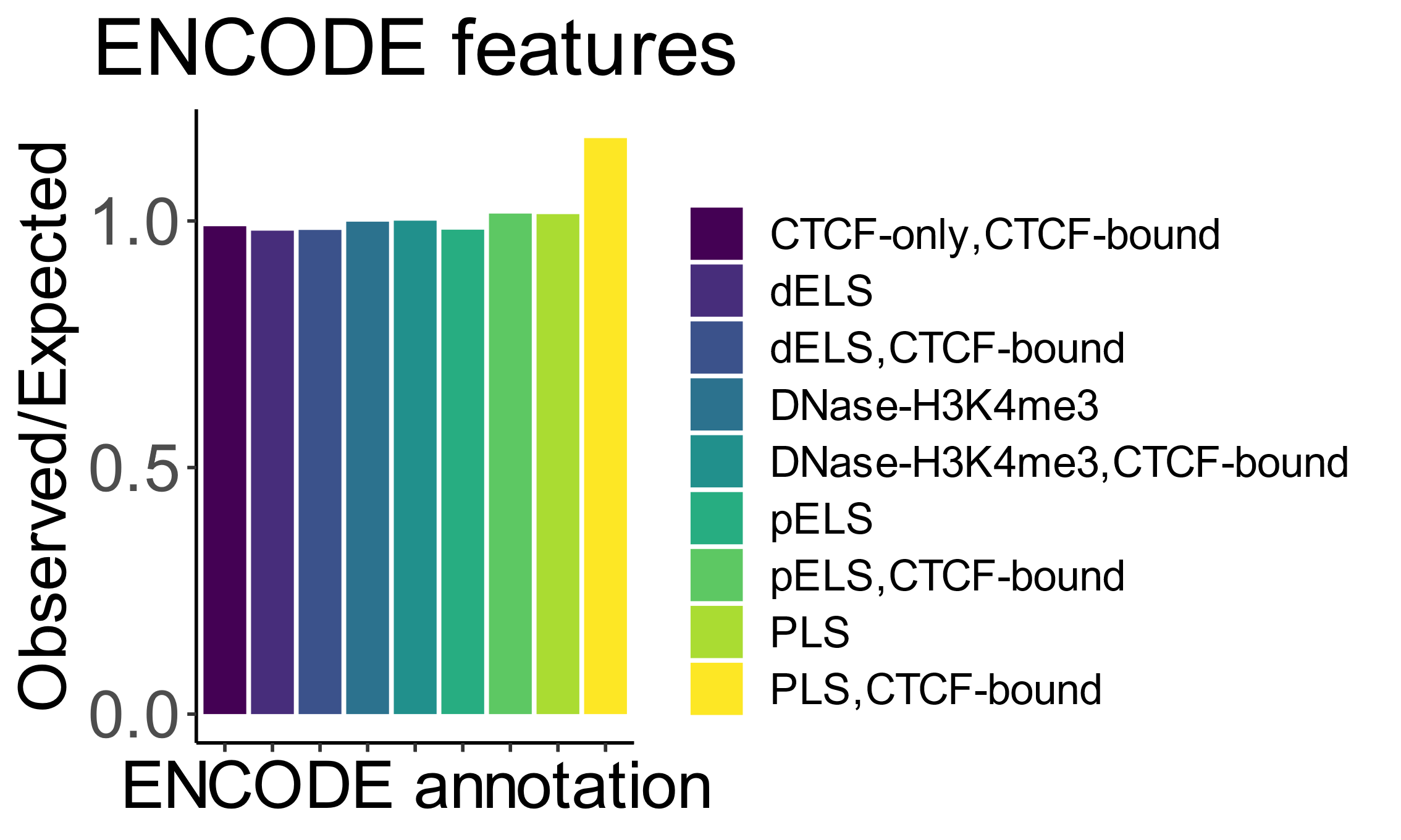

**Supplementary Figure 8. Roulette performance at different DNA regions, as annotated by ENCODE.**

Observed to expected ratio of rare SNVs at different ENCODE annotations. PLS stands for promoters, ELS for enhancers.

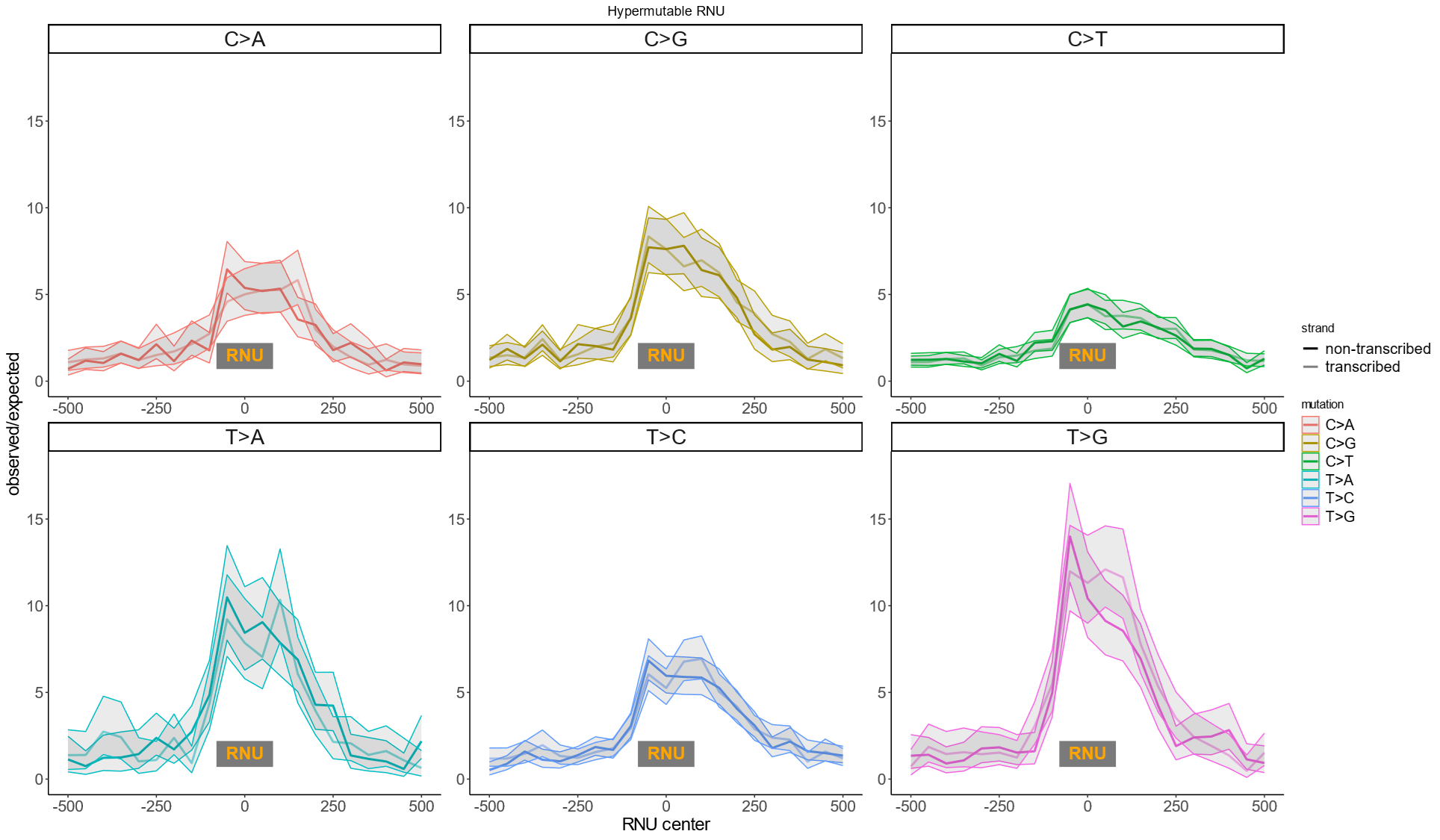

**Supplementary Figure 9. Mutation rate around RNU genes.** Shaded area is Poisson confidence intervals.

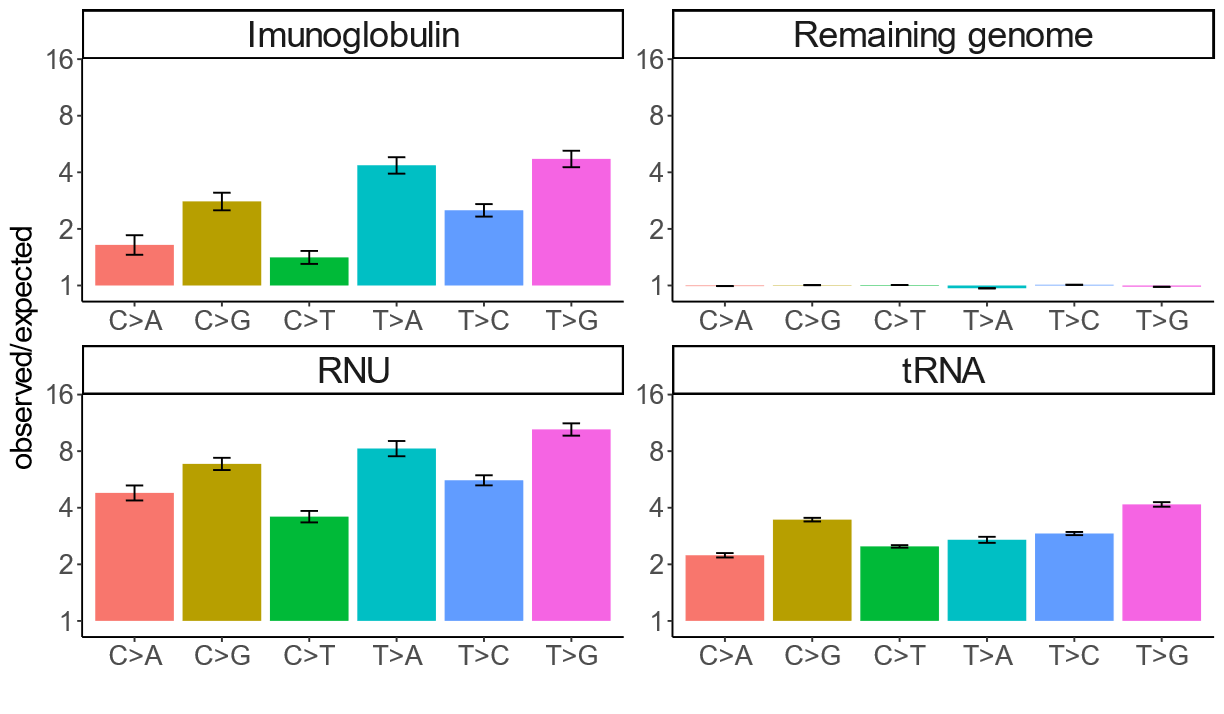

**Supplementary Figure 10. Mutation rate elevation in hypermutable gene classes decomposed by mutation type.** We quantified the mutation rate elevation for each mutation type in order to visualize changes to the mutation spectrum in gene classes with elevated SNV counts.

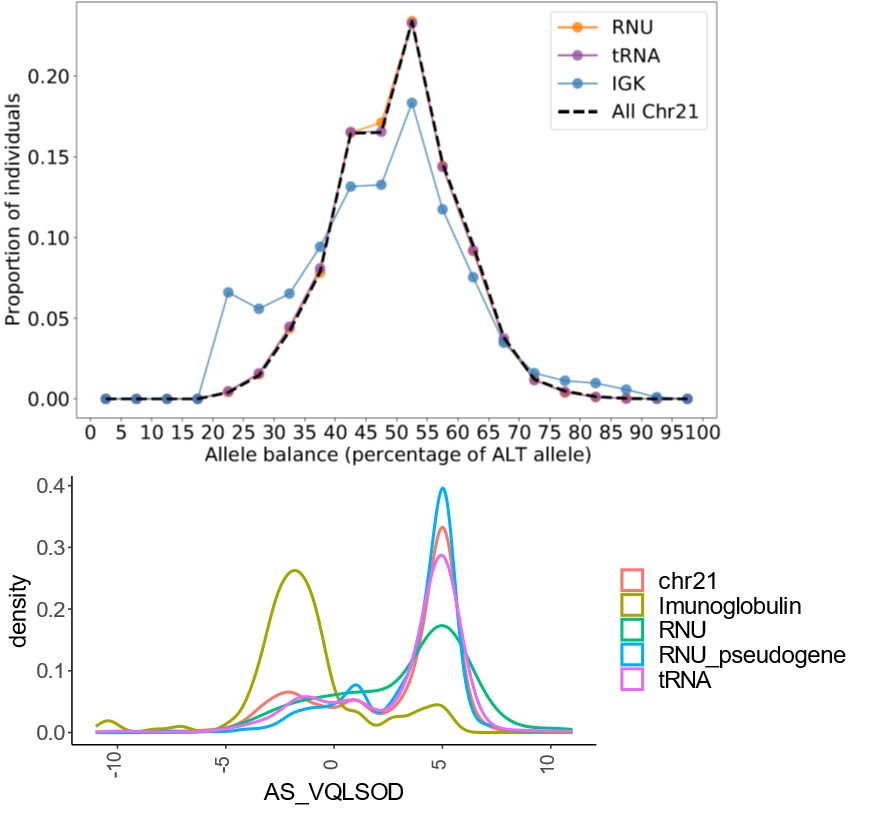

**Supplementary Figure 11. Quality metrics for hypermutable regions.** a) The allelic balance within RNU and tRNA genes match the control curve for all sites on chromosome 21, while IGK genes deviate substantially from the background. b) AS_VQSLOD which is the main metric for the quality of the variant is dramatically decreased for IGK.

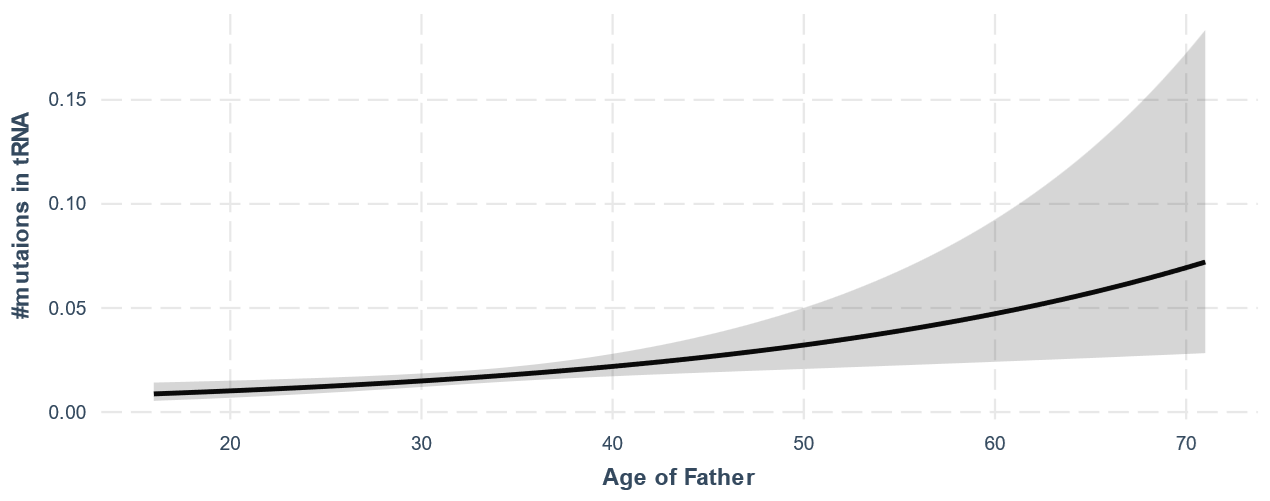

**Supplementary Figure 12. Parental age effect**. The best fit for the relationship between the number of mutations occurring in tRNA sites and the parental age. The fit is for the exponential dependency on age using Poisson regression. While only 104 *de novo* mutations occurred in tRNA genes, the association between parental age and the number of mutations is highly significant (p=9.9*10^-5^, results remain significant for the linear relationship between age and mutation count: p=5.6*10^-3^). Only 18 mutations happened in RNU genes making analysis of the age effect unreliable.

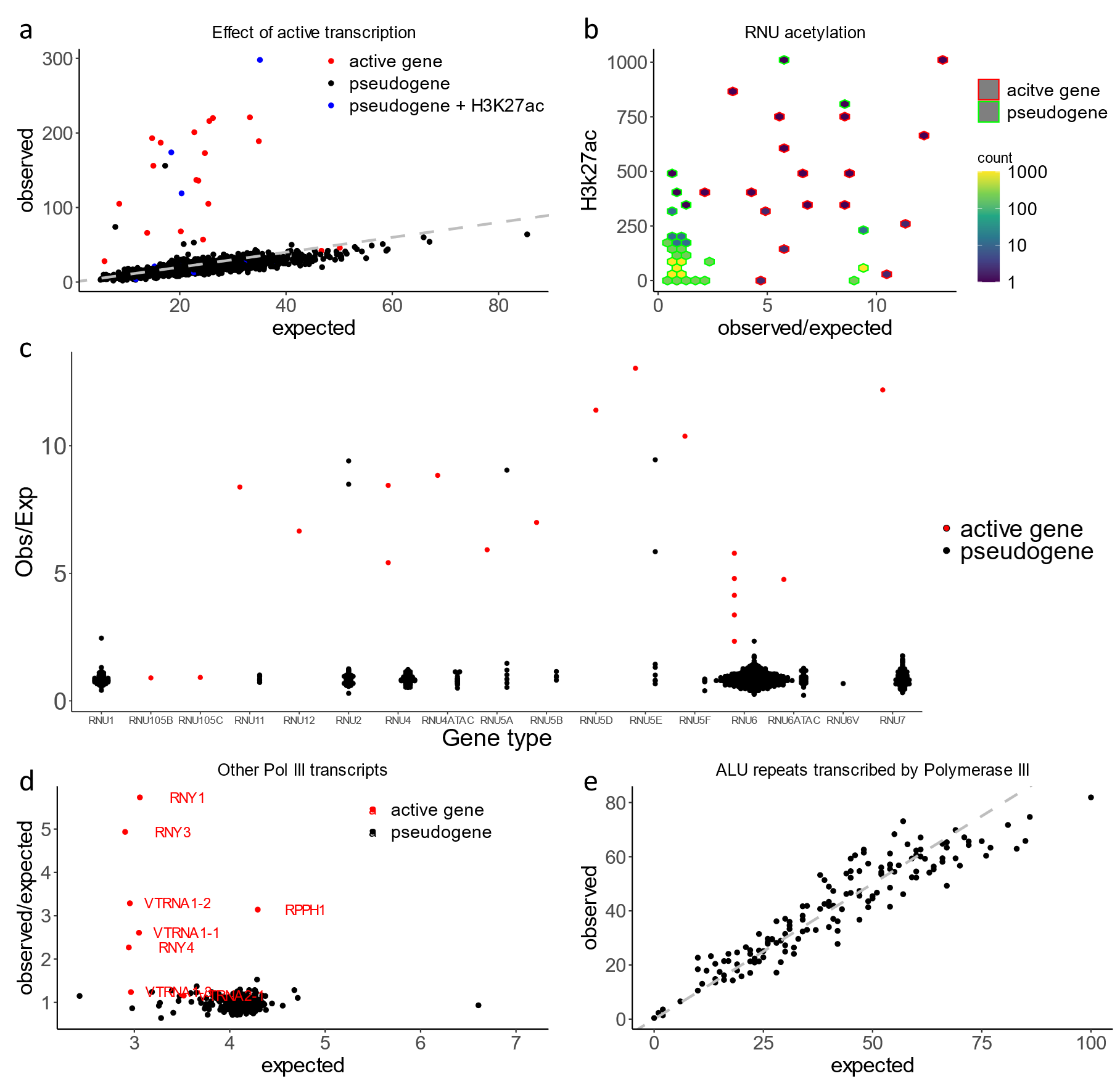
**Supplementary Figure 13. Effect of polymerase III transcription**

a,b) The effect of acetylation and pseudo/active gene stratification (according to the HUGO annotation) on the SNV number in RNU genes. b) While the vast majority of RNU pseudogenes do not overlap with H3k27ac peaks, four out of five RNU pseudogenes with high mutation rate overlap H3k27ac. c) Active transcription increases mutation rate across different classes of RNU genes. d) Other classes of Pol III transcript, represented by a small number of genes also have elevated mutation rate e) The density of rare SNVs is in line with the Roulette predictions in ALU elements that have been predicted to be transcribed by polymerase III.

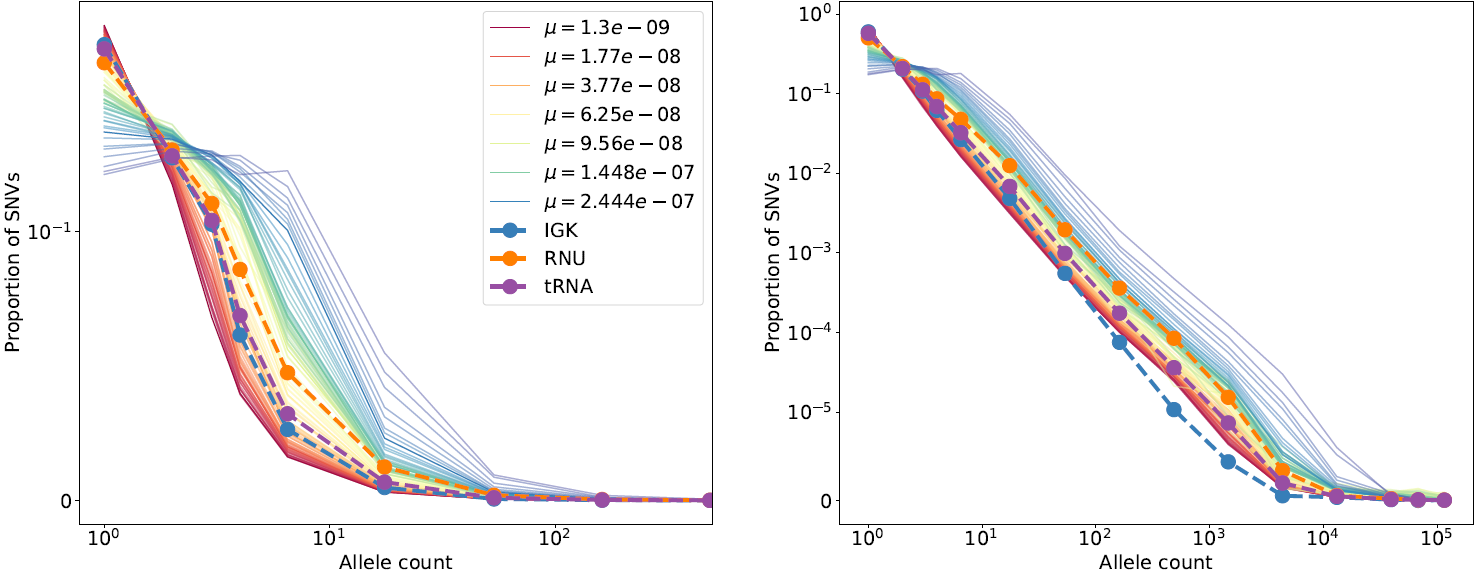

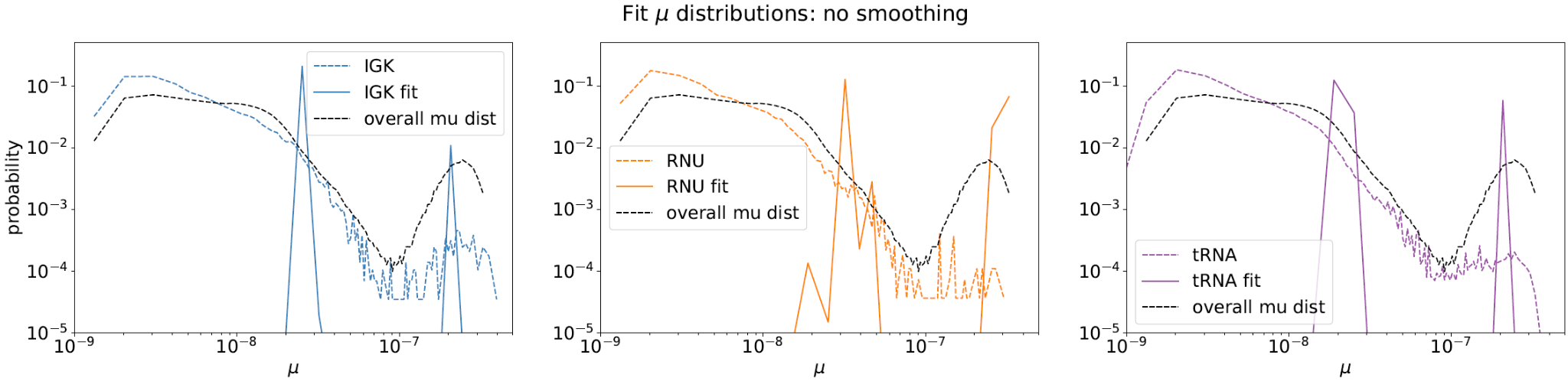

**Supplementary Figure 14. Site frequency spectra (SFS) for sites with different mutation rates and for different gene categories.** Rare variants on the top left and full SFS to the top right. On the bottom the mutation rate distributions for observed SNVs for different gene categories was estimated by fitting the SFS in these genes as a mixture of SFS shapes observed in Roulette bins. In contrast to Figure 4 c, d we did not use genome-wide mutation rate distribution as a prior.

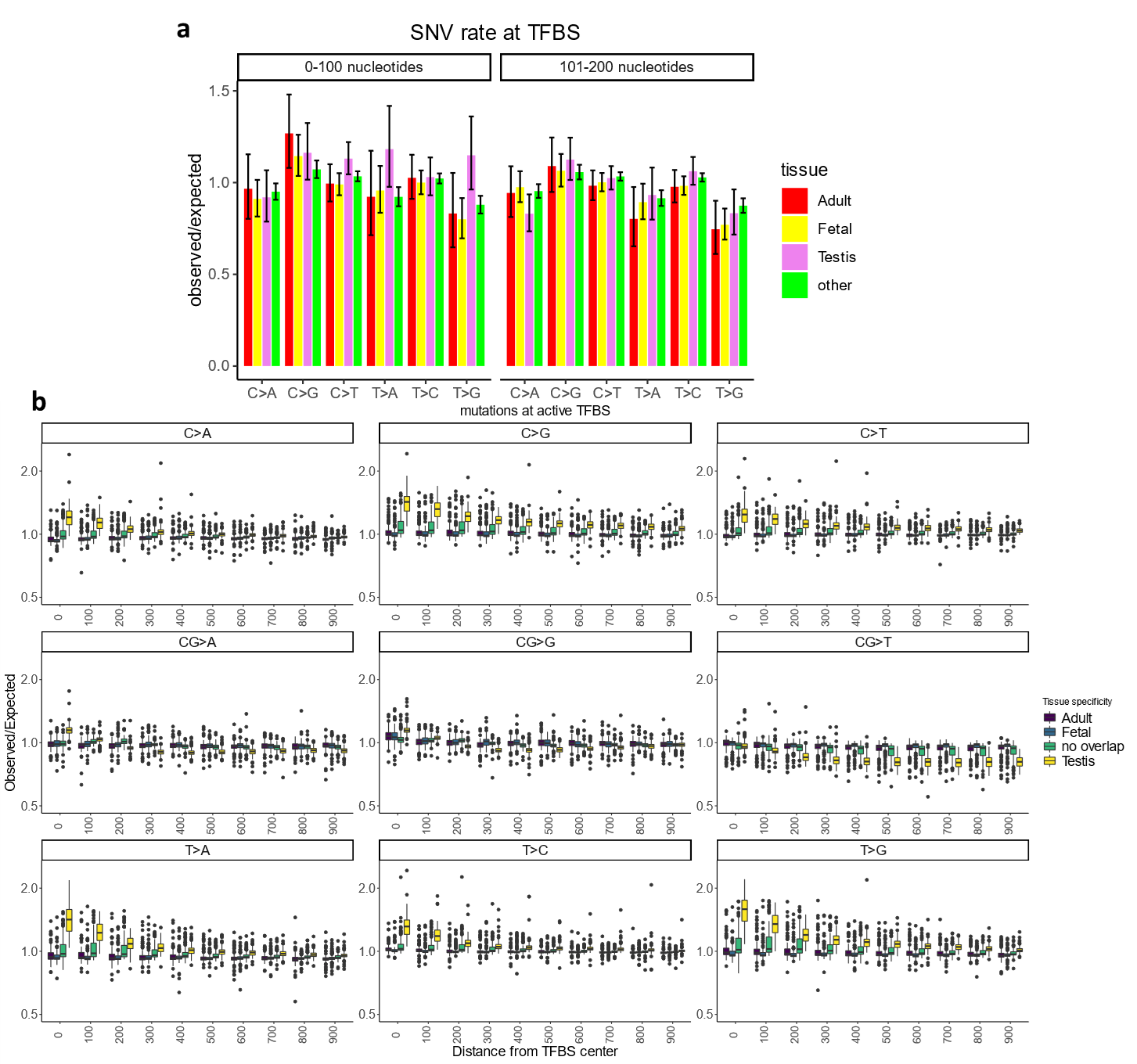

**Supplementary Figure 15. Mutation rate is accelerated at TFBS active in testis**

a) Observed to expected ratio of *de novo* mutations at TFBS active in different tissues. Distance from the TFBS center is shown on top. b) Observed to expected ratio of rare SNVs at TFBS active in different tissues. Panels are stratified by mutation type.

**Supplementary Figure 16. Mutation rate is accelerated at TFBS active and overlapping multiple promoters.** We compared observed/expected mutation rate for non- CpG mutations overlapping DHS in testis. These TFBS have higher mutation rate if they also overlap multiple promoters (light yellow) instead of a single promoter (dark yellow).
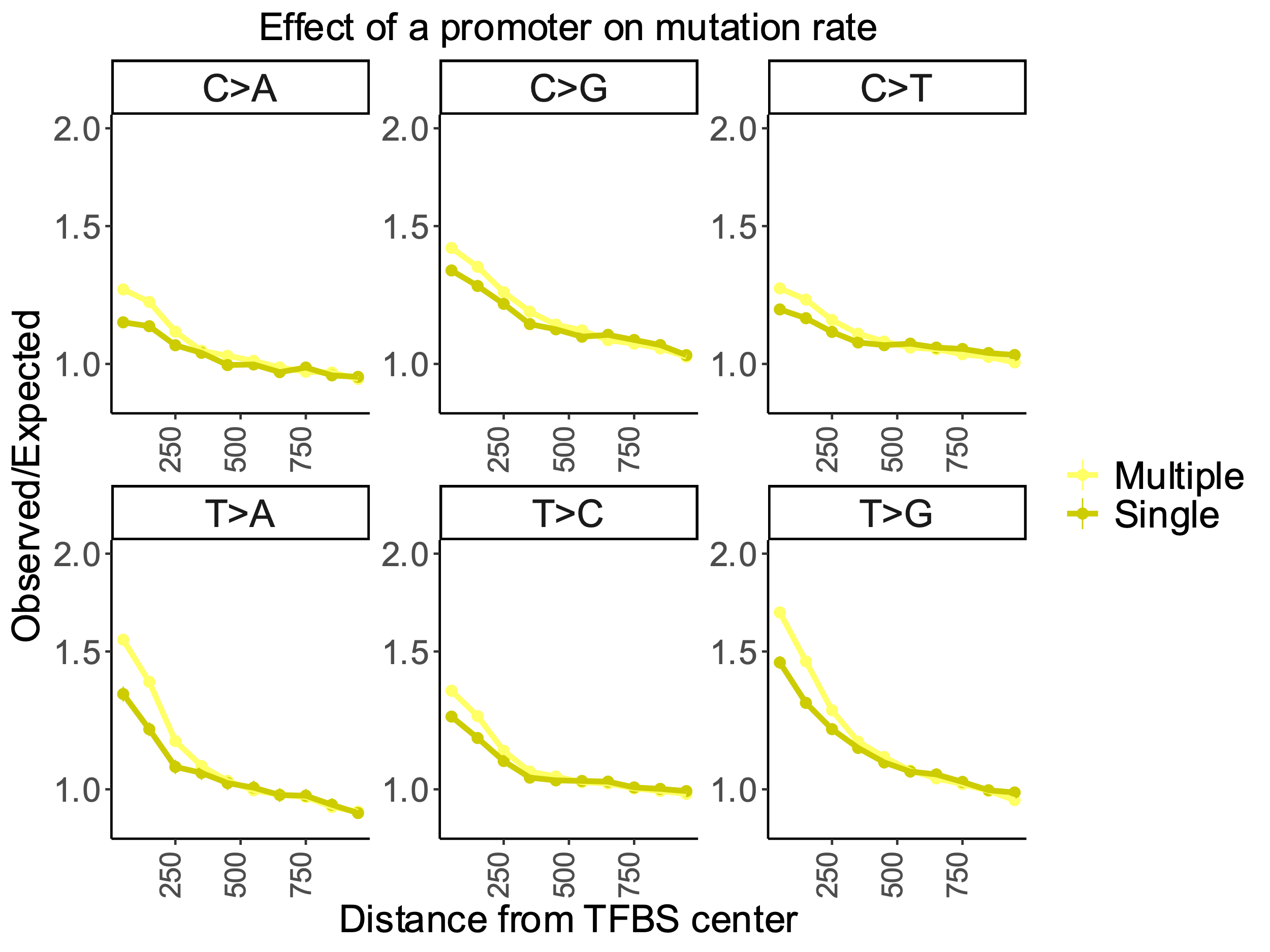

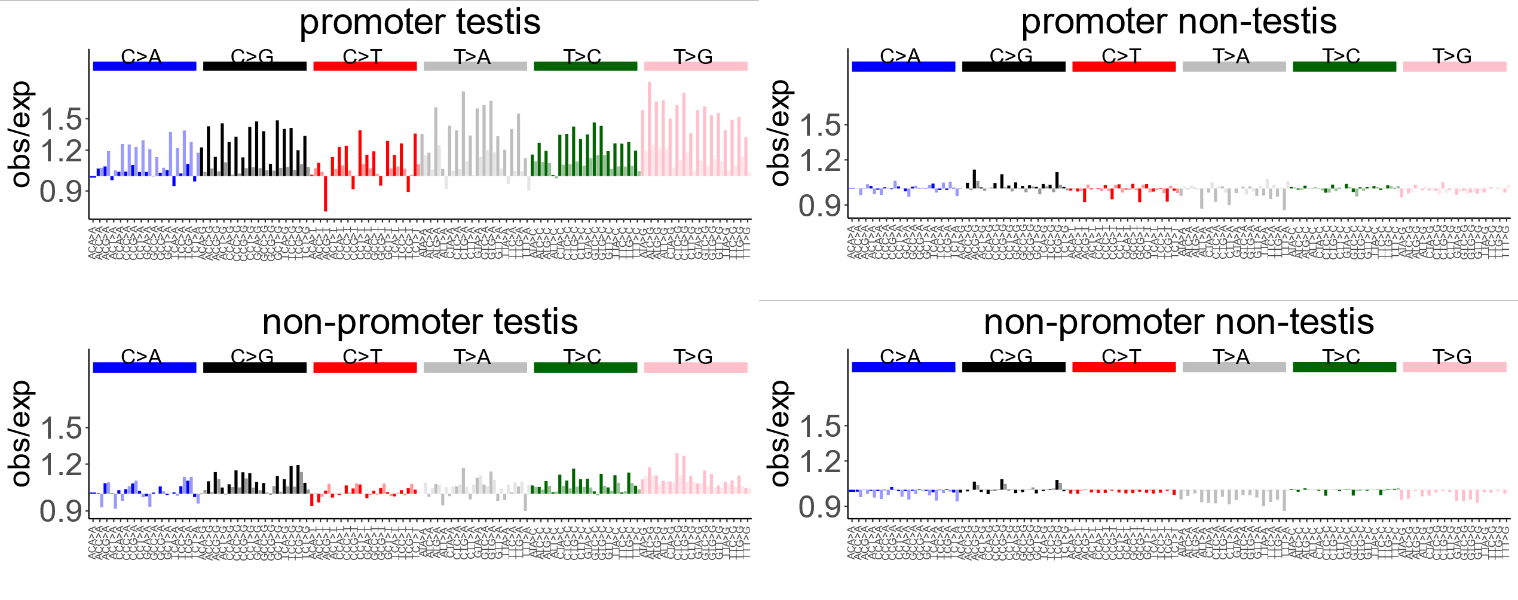

**Supplementary Figure 17. Correction of mutation rate at TFBS**

Panels show the observed to expected density of rare SNVs at TFBS active in Testis or in other tissues as well as whether TFBS do or do not overlap a promoter. Bright colors are reflecting deviation from the model before the correction for higher mutability at TFBS, pale colors correspond for corrected values.

**Supplementary Figure 18.** **UV-induced mutations in melanoma.** We analyzed TFBS overlapping DHS of foreskin melanocytes and measured the rate of TCC>T mutations (major UV-induced mutation type). Mutation rates in melanoma samples (ICGC data) differ between TFBS sites overlapping and not overlapping promoters. Results are in line with Mao et al, Nature Comms. 2018
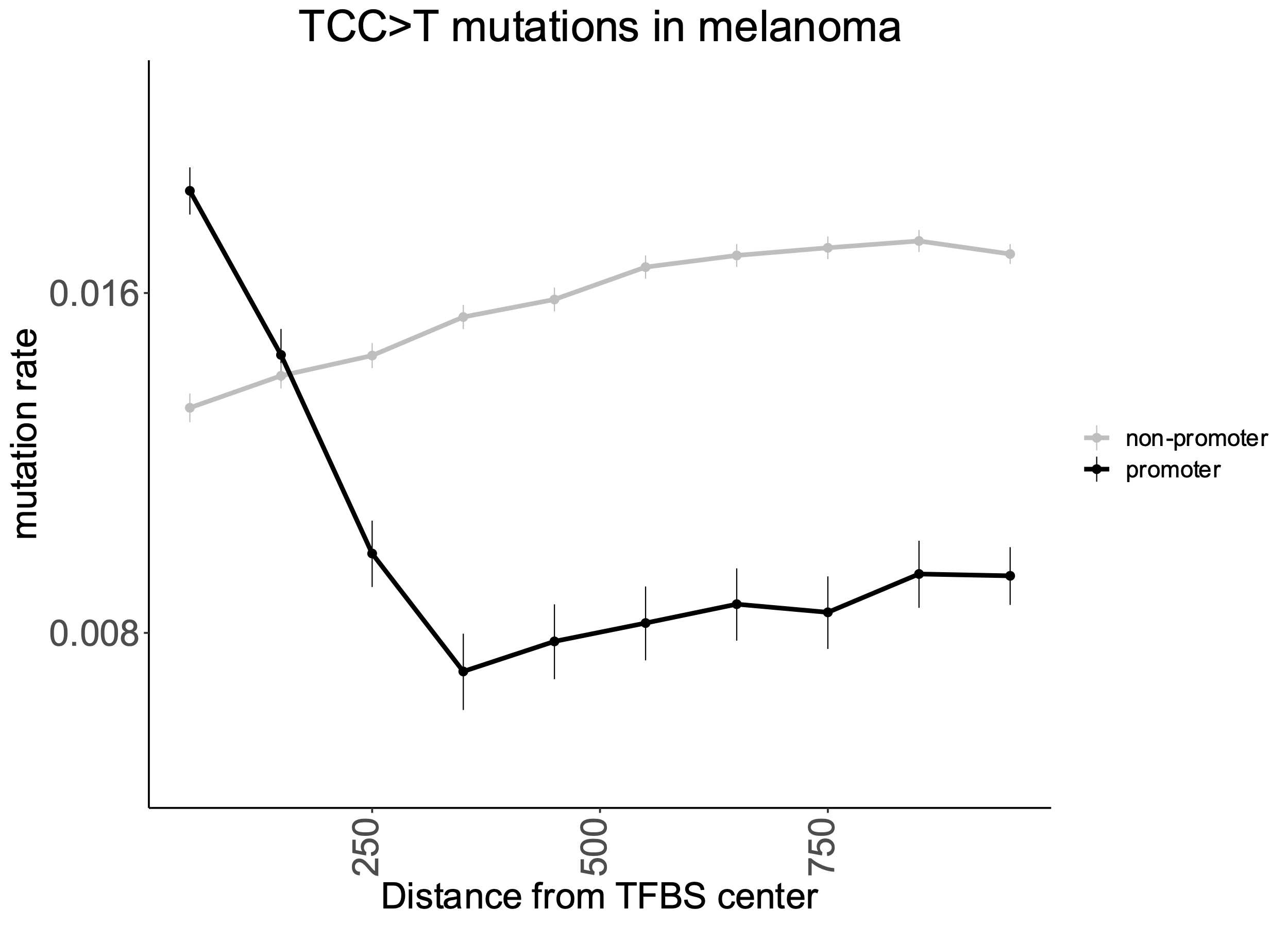

**Supplementary Figure 19. Effect of surrounding nucleotides is varying between the coding strand, the non-coding strand within genes and intergenic regions**
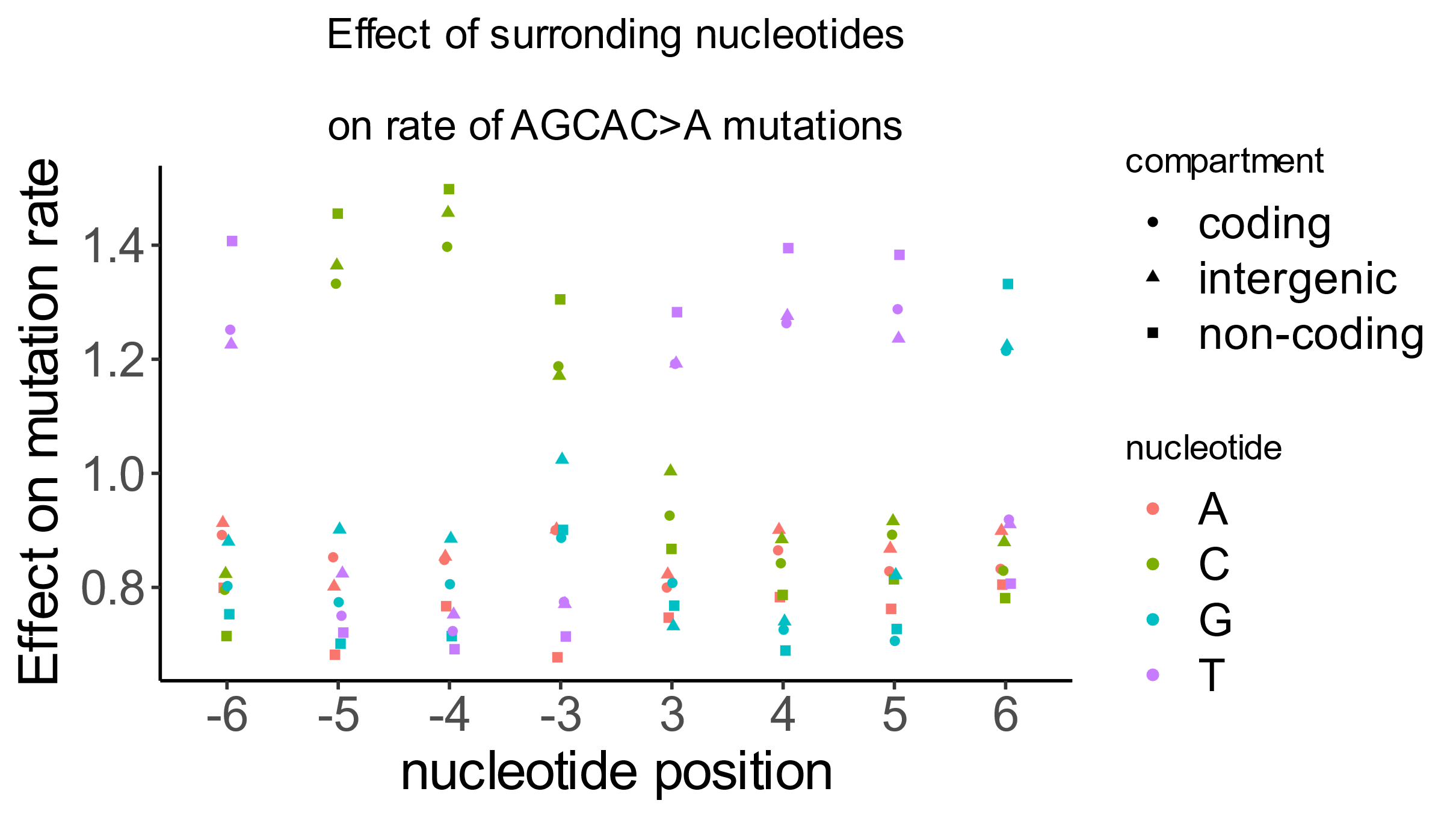

The figure shows the effect of nucleotides beyond the pentamer on the rate of AGCAC>A mutations. The effect differs between genes and intergenic regions and is strand-dependent within genes. The AGCAC>A mutation type is shown as an example.

**Supplementary Figure 20. Filtering pentameric contexts with abnormal patterns of site frequency spectra**
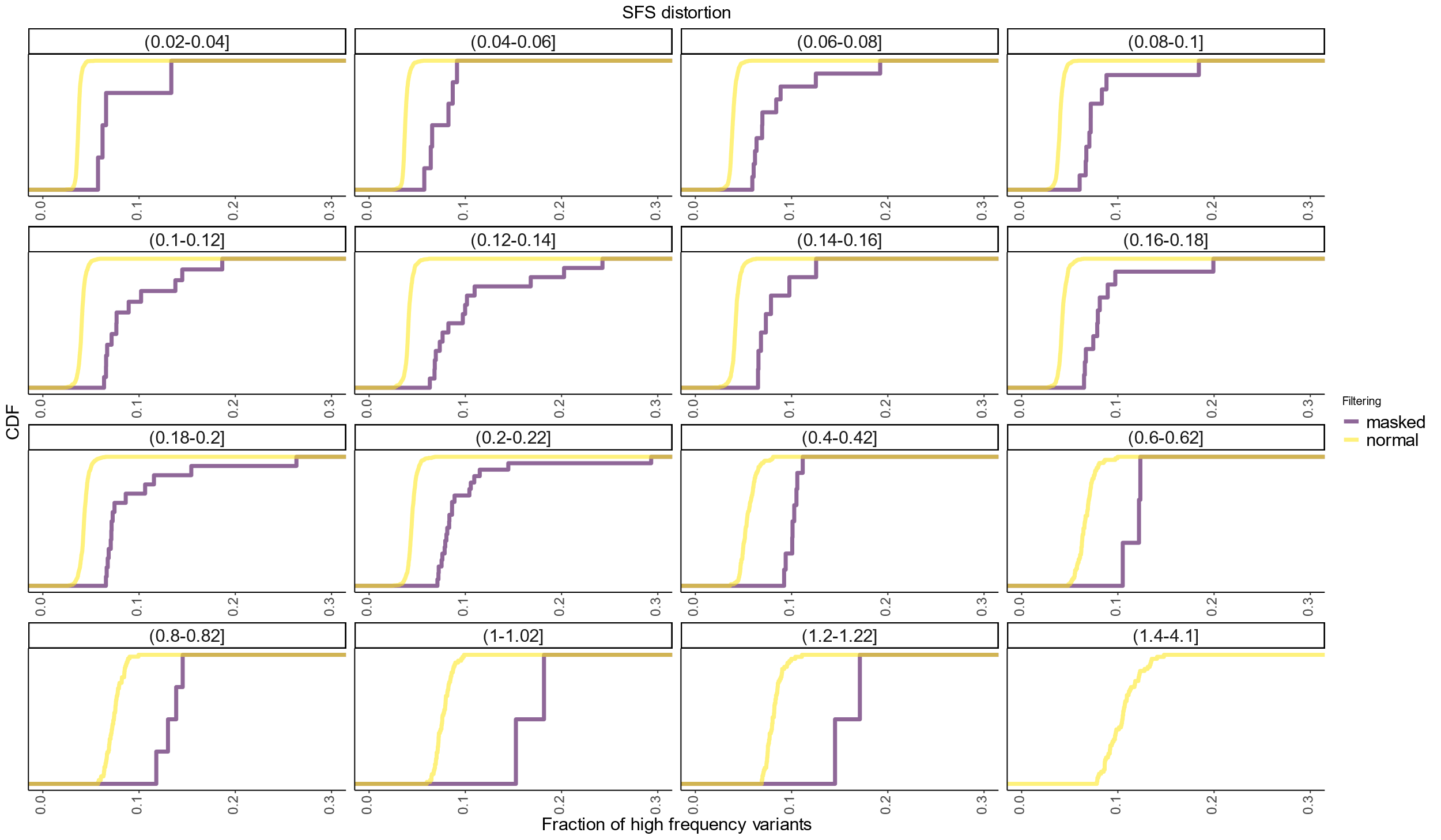

Site frequency spectrum (SFS) is dependent on mutation rate for rare SNVs. For each pair of pentamer and predicted mutation rate (shown on top of the panels) we calculated the fraction of high frequency variants (MAF >0.005 and MAF <=0.2). We masked the context if it has a proportion of high frequency variants exceeding mean for the same mutation rate multiplied by 1.5. We show empirical cumulative distributions for the portion of high frequency variants for masked (purple) and non-masked pentamers.

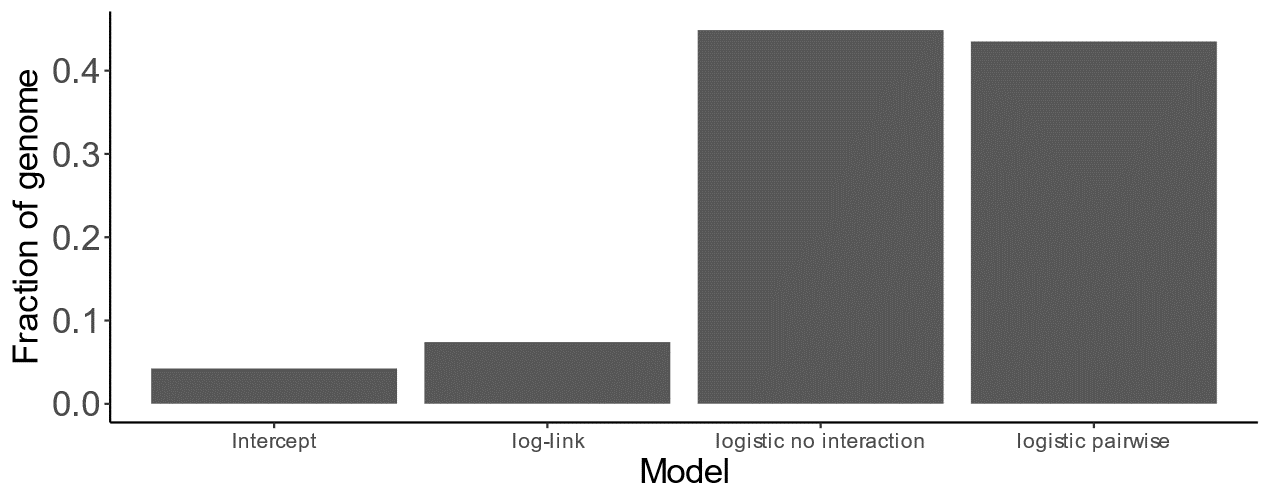

**Supplementary Figure 21. Fraction of the genome where each of four tested models fits best.** We selected the best of four models on a 50% hold out test for each genomic compartment.

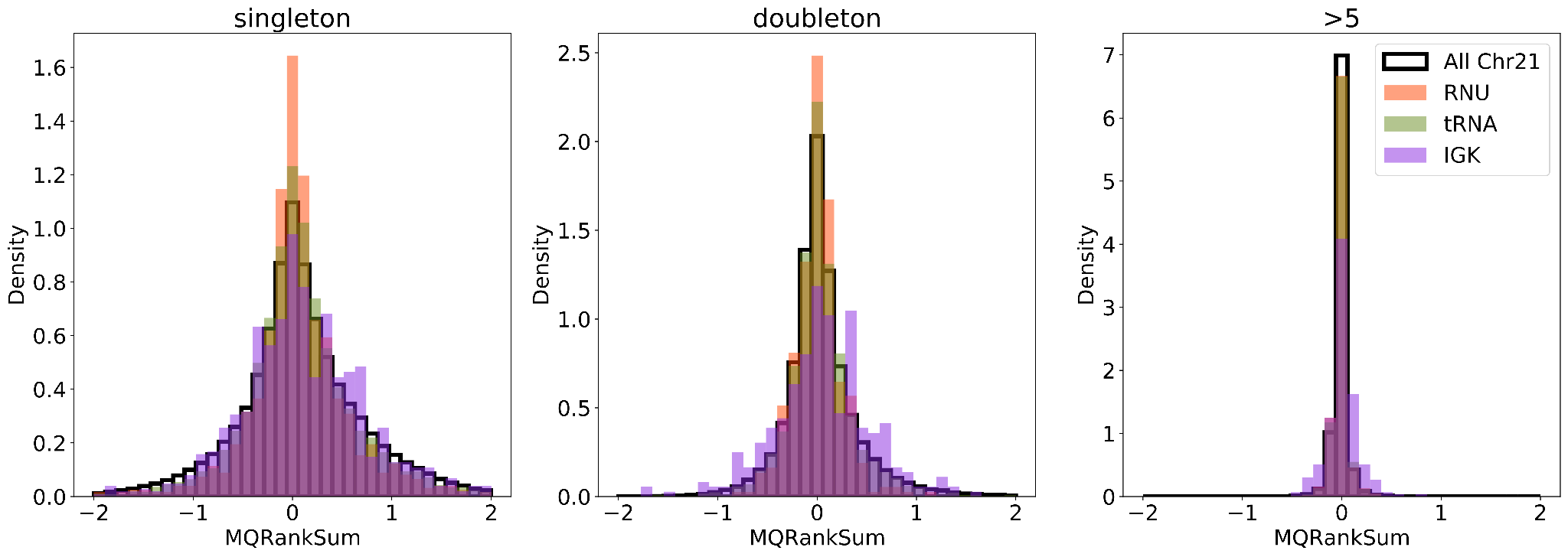

**Supplementary Figure 22. Distributions of mapping quality scores for apparent hypermutable gene classes.** We show the distribution of MQRankSum scores for SNVs in RNU, tRNA, and IGK genes compared to the background distribution from chromosome 21. Distributions were computed from the 1K genomes subset of gnomAD v3.

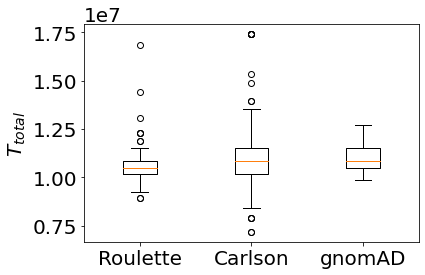

**Supplementary Figure 23. Distribution of** $T_{total}$ **values fit for each model in the SNV-conditional procedure.**

| **Model** | **Training Set** | **Outcome Variable** | **Regression**  **Type** | **Context** | **Regional** | **Covariates** |
| --- | --- | --- | --- | --- | --- | --- |
| Roulette | gnomAD v3,  intergenic sites | Polymorphic Sites | Logistic | Pentamer + surrounding nucleotides | 50kb window | -methylation  -transcription asymmetry  -more features |
| Carlson | BRIDGES study, all singletons | Presence of Singletons | Logistic | Heptamer | None | -methylation  -more features |
| gnomAD | gnomAD v2,  noncoding region | Proportion observed for context | Proportion | Trimer | None | -methylation |

Other covariates by Carlson:

- Replication Timing
- Recombination Rate
- Lamin B1 domains
- DNase hypersensitivity sites
- In Exon
- CpG Island
- % GC Content

**Supplementary Table 1. Overview of Different Mutation Rate Models**

Summary of the three mutational models.

| Method | Roulette AUC  (Precision-Recall curve) | Roulette genes | gnomAD AUC | gnomAD genes | Roulette auc - gnomAD auc |
| --- | --- | --- | --- | --- | --- |
| shet | 0.298  [0.273, 0.324] | GRIN2A  AUTS2  CSNK2A1  AP2S1  GLI3  ZIC1  TSC2 | 0.289  [0.267, 0.316] | PTHLH | 0.007  [0.002, 0.014] |
| LOEUF | 0.291  [0.268, 0.317] | FGF10  FOXF1  PITX2  PUF60  WT1 | 0.287  [0.262, 0.312] | CAMK2G  KCNC1 | 0.004  [-0.002, 0.01] |

**Supplementary Table 2. Performance of shet and LOEUF based on Developmental Disease Genotype to Phenotype Database (DDG2P) genes with highest confidence value**

We re-estimate shet and LOEUF using gnomAD.v2.1.1 whole exomes, under both Roulette rates and gnomAD rates. We then use a set of developmental disease genes to test the performance of the two shet values. Due to data imbalance, the AUC for Precision-Recall curve was used. For the AUC, we bootstrap all of the genes and show the 95% confidence interval.

**Supplementary Table 3. Contexts with high fraction of high frequency polymorphisms**

| Mutation rate category | Context | Enrichment of high frequency variants |
| --- | --- | --- |
| (0.02-0.04] | TATAT_G | 1.637619 |
| (0.02-0.04] | TCTCA_A | 3.297963 |
| (0.02-0.04] | TTTTA_A | 1.634093 |
| (0.02-0.04] | TTTTT_A | 1.516249 |
| (0.04-0.06] | CATCC_G | 1.656761 |
| (0.04-0.06] | TATAT_G | 2.094389 |
| (0.04-0.06] | TGTGA_A | 2.176461 |
| (0.04-0.06] | TTTTA_A | 2.190425 |
| (0.04-0.06] | TTTTT_A | 1.632656 |
| (0.06-0.08] | AACTG_T | 1.55859 |
| (0.06-0.08] | AATAT_A | 1.79443 |
| (0.06-0.08] | GGTGT_G | 1.615888 |
| (0.06-0.08] | TGTGG_G | 1.725487 |
| (0.06-0.08] | TTTTA_A | 3.052376 |
| (0.06-0.08] | TTTTG_G | 2.211894 |
| (0.06-0.08] | TTTTT_A | 1.610359 |
| (0.08-0.1] | AATAA_A | 1.761103 |
| (0.08-0.1] | AGTGT_A | 1.680997 |
| (0.08-0.1] | GGTGT_G | 2.105791 |
| (0.08-0.1] | TGTGG_G | 2.058173 |
| (0.08-0.1] | TTTTA_A | 4.006997 |
| (0.08-0.1] | TTTTT_A | 1.7158 |
| (0.08-0.1] | TTTTT_G | 1.739209 |
| (0.1-0.12] | AATAA_A | 2.080842 |
| (0.1-0.12] | AATAT_A | 2.429194 |
| (0.1-0.12] | ATTTT_A | 1.716421 |
| (0.1-0.12] | GGTGT_G | 3.552388 |
| (0.1-0.12] | TGTTG_G | 2.066228 |
| (0.1-0.12] | TGTTT_G | 1.556219 |
| (0.1-0.12] | TTTTA_A | 4.199835 |
| (0.1-0.12] | TTTTG_G | 3.247131 |
| (0.12-0.14] | AATAA_A | 2.464163 |
| (0.12-0.14] | AATAT_A | 2.205127 |
| (0.12-0.14] | ATTTT_A | 2.166376 |
| (0.12-0.14] | CCCCT_G | 1.54065 |
| (0.12-0.14] | TTTAA_A | 1.550984 |
| (0.12-0.14] | TTTGG_G | 1.54998 |
| (0.12-0.14] | TTTTA_A | 4.078965 |
| (0.12-0.14] | TTTTG_G | 3.497853 |
| (0.14-0.16] | AATAA_A | 2.840911 |
| (0.14-0.16] | ATTTT_A | 2.109861 |
| (0.14-0.16] | CCCCA_G | 1.512262 |
| (0.14-0.16] | CCCCT_G | 1.682919 |
| (0.14-0.16] | GTTTT_G | 1.5084 |
| (0.14-0.16] | TTTAA_A | 1.741826 |
| (0.16-0.18] | AATAA_A | 2.134233 |
| (0.16-0.18] | ATTTT_A | 1.722081 |
| (0.16-0.18] | CCCCA_G | 1.575511 |
| (0.16-0.18] | CCCCT_G | 1.898158 |
| (0.16-0.18] | TTTAA_A | 1.710133 |
| (0.16-0.18] | TTTCC_C | 1.504895 |
| (0.18-0.2] | AATAA_A | 2.298695 |
| (0.18-0.2] | AATAT_A | 5.107503 |
| (0.18-0.2] | CACAA_A | 1.539471 |
| (0.18-0.2] | CCCCA_A | 1.564379 |
| (0.18-0.2] | CCCCA_G | 1.672006 |
| (0.18-0.2] | CCCCC_G | 1.606482 |
| (0.18-0.2] | CCCCT_G | 2.08087 |
| (0.18-0.2] | GGTGT_G | 3.332767 |
| (0.18-0.2] | GTTTT_G | 1.783016 |
| (0.18-0.2] | TTTAA_A | 1.628809 |
| (0.18-0.2] | TTTGG_G | 2.500418 |
| (0.2-0.22] | AACCA_A | 1.730352 |
| (0.2-0.22] | AACGA_T | 1.548946 |
| (0.2-0.22] | AATAA_A | 2.038241 |
| (0.2-0.22] | AATAT_A | 4.591603 |
| (0.2-0.22] | AATTT_A | 1.794179 |
| (0.2-0.22] | CACAA_A | 1.962771 |
| (0.2-0.22] | CACGC_T | 1.503114 |
| (0.2-0.22] | CCCAA_A | 1.908264 |
| (0.2-0.22] | CCCAC_G | 1.504297 |
| (0.2-0.22] | CCCCA_A | 1.518981 |
| (0.2-0.22] | CCCCC_A | 1.545511 |
| (0.2-0.22] | CCCCC_G | 1.730979 |
| (0.2-0.22] | CCCCT_G | 1.881438 |
| (0.2-0.22] | CTCAA_A | 2.034548 |
| (0.2-0.22] | GACGC_T | 1.573817 |
| (0.2-0.22] | GATAT_G | 1.60293 |
| (0.2-0.22] | GCCCC_A | 1.606621 |
| (0.2-0.22] | GGTTT_G | 2.029864 |
| (0.2-0.22] | TGTTT_G | 1.50068 |
| (0.2-0.22] | TTTAA_A | 2.137843 |
| (0.2-0.22] | TTTAT_A | 1.597595 |
| (0.2-0.22] | TTTCC_C | 1.676408 |
| (0.2-0.22] | TTTGG_G | 2.10929 |
| (1.4-4.1] | AATTT_A | 2.872091 |
| (1.4-4.1] | TTTAA_A | 2.928568 |
